# Pharmacological and genetic activation of cAMP synthesis disrupts cholesterol utilization in *Mycobacterium tuberculosis*

**DOI:** 10.1101/2021.08.03.454881

**Authors:** Kaley M. Wilburn, Christine R. Montague, Bo Qin, Ashley K. Woods, Melissa S. Love, Case W. McNamara, Peter G. Schultz, Teresa L. Southard, Lu Huang, H. Michael Petrassi, Brian C. VanderVen

## Abstract

There is a growing appreciation for the idea that bacterial utilization of host-derived lipids, including cholesterol, supports *Mycobacterium tuberculosis* (Mtb) pathogenesis. This has generated interest in identifying novel antibiotics that can disrupt cholesterol utilization by Mtb *in vivo*. Here we identify a novel small molecule agonist (V-59) of the Mtb adenylyl cyclase Rv1625c, which stimulates 3’, 5’-cyclic adenosine monophosphate (cAMP) synthesis and inhibits cholesterol utilization by Mtb. Similarly, using a complementary genetic approach that induces bacterial cAMP synthesis independent of Rv1625c, we demonstrate that inducing cAMP synthesis is sufficient to inhibit cholesterol utilization in Mtb. Although the physiological roles of individual adenylyl cyclase enzymes in Mtb are largely unknown, here we demonstrate that the transmembrane region of Rv1625c is required for cholesterol metabolism. Finally, in this work the pharmacokinetic properties of Rv1625c agonists are optimized, producing an orally-available Rv1625c agonist that impairs Mtb pathogenesis in infected mice. Collectively, this work demonstrates a novel role for Rv1625c and cAMP signaling in controlling cholesterol metabolism in Mtb and establishes that cAMP signaling can be pharmacologically manipulated for the development of new antibiotic strategies.

**Author Summary:** The recalcitrance of *Mycobacterium tuberculosis* (Mtb) to conventional antibiotics has created a need to identify novel pharmacological mechanisms to inhibit Mtb pathogenesis. There is a growing understanding of the metabolic adaptations Mtb adopts during infection to support its survival and pathogenesis. This has generated interest in identifying small molecule compounds that effectively inhibit these *in vivo* metabolic adaptations, while overcoming challenges like poor pharmacokinetic properties or redundancy in target pathways. The Mtb cholesterol utilization pathway has repeatedly been speculated to be a desirable antibiotic target, but compounds that successfully inhibit this complex pathway and are suitable for use *in vivo* are lacking. Here, we establish that stimulating cAMP synthesis in Mtb is a mechanism that is sufficient to block cholesterol utilization by the bacterium, preventing the release of key metabolic intermediates that are derived from breakdown of the cholesterol molecule. For the first time, we identify small molecule agonists of the Mtb adenylyl cyclase Rv1625c that have promising pharmacological properties and are suitable for use during *in vivo* studies. These Rv1625c agonists increase cAMP synthesis, inhibit cholesterol utilization by Mtb, and disrupt Mtb pathogenesis in mouse models of chronic infection.

## Introduction

Tuberculosis (TB) remains a prevalent infectious disease worldwide that claims ∼1.4 million lives and afflicts ∼10 million new individuals annually (*1*). TB is caused by *Mycobacterium tuberculosis* (Mtb), and it is an ongoing challenge to identify antibiotics with novel bacterial targets that can shorten treatment, limit side-effects, and reduce disease relapse. An important aspect of Mtb pathogenesis is that the bacterium persists in the human lung within lipid-rich phagocytes and/or tissue lesions while promoting pathology that is required for dissemination and transmission (*2*). Mtb primarily lives within macrophages and stimulates the formation of lipid-loaded cells (*3, 4*), but the bacterium can also survive in the acellular core of necrotic granulomas that are rich in cholesterol, cholesterol ester, and triacylglycerol (*2, 5*). It is generally understood that Mtb utilizes host-derived lipids, including cholesterol, as key nutrients to survive during persistent infection (*6*). Mtb completely degrades cholesterol into two- and three-carbon intermediates that are metabolized for energy production or serve as biosynthetic precursors of cell wall or virulence lipids (*6*). In animal models, Mtb requires cholesterol metabolism to maintain optimal chronic lung infection (*7–11*) and cholesterol utilization was recently found to belong to a set of “core virulence functions” required for Mtb survival *in vivo* across a genetically diverse panel of mice (*12*). Furthermore, it was recently demonstrated that a multi-drug resistant strain of Mtb is more dependent on cholesterol for growth than an H37Rv reference strain (*13*). Thus, the cholesterol metabolic pathway in Mtb represents a novel, genetically validated target for drug discovery. However, tools to pharmacologically inhibit this pathway during infection *in vivo* have yet to be developed.

Signaling through the universal second-messenger 3’,5’-cyclic adenosine monophosphate (cAMP) has long been studied in a variety of prokaryotic and eukaryotic systems. In pathogenic bacteria, cAMP is essential in regulating functions such as carbon metabolism, virulence gene expression, biofilm formation, drug tolerance, and manipulation of host cell signaling (*14–16*). How cAMP signaling regulates Mtb physiology during infection is not well understood, partly due to the limited tools available for investigating this and the myriad of pathway components present in Mtb. The Mtb genome encodes an unusually large repertoire of at least ten biochemically active class III adenylyl cyclase (AC) enzymes, which catalyze the intramolecular cyclization of ATP to form cAMP when activated. These ACs are structurally diverse, and the majority of these proteins are composed of a catalytic domain along with other accessory domains, which are thought to participate in regulatory or effector functions (*17*). Studies using recombinant expression systems have proposed environmental stimuli (e.g. pH, fatty acids, or HCO_3_^-^/CO_2_) for five Mtb ACs (*18–22*). Additionally, Mtb possesses twelve predicted downstream cAMP-binding effector proteins, only four of which have been functionally characterized (*23–28*). Thus, our understanding of how individual ACs and downstream cAMP-dependent effector proteins regulate specific aspects of Mtb physiology is extremely limited. To date, no individual AC enzyme has been directly linked to the regulation of a specific biological or metabolic process in Mtb.

We previously identified a series of compounds that inhibit Mtb growth in macrophages and in a cholesterol media (*29*). The activity of a subset of these compounds was dependent on the AC Rv1625c, and compound treatment increased cAMP production in Mtb (*29*). The Rv1625c protein is composed of at least four structural elements: an N-terminal cytoplasmic tail, a six-helical transmembrane domain, a cytoplasmic helical domain, and a C-terminal cyclase domain. Based on its topology and sequence homology, Rv1625c is comparable to ‘one-half’ of a mammalian membrane-associated AC (*30*). Rv1625c forms a homodimer to generate two active sites composed of complementary residues, and conserved active site residues as well as the cytoplasmic tail and helical domain have been linked to its catalytic activity (*31–33*). Although it has been proposed that Rv1625c may be activated by binding HCO_3_^-^/CO_2_ or lipophilic ligands, it remains unclear what the native role of Rv1625c is in Mtb during infection (*19, 34, 35*). The possibility that we had identified chemical tools comparable to forskolin in the Mtb system led us to investigate the mechanism of these Rv1625c-dependent compounds and their impact on Mtb carbon metabolism and pathogenesis. We were especially motivated to test the hypothesis that activating cAMP synthesis in Mtb through an Rv1625c agonist could disable cholesterol utilization and undermine Mtb persistence during infection in mice.

To carry out these studies, we re-examined our previously identified screening hits for Rv1625c-dependent compounds with favorable pharmacokinetic properties. From these, we selected a potent compound (V-59) that permitted both *in vitro* and *in vivo* studies to examine the impact that chemically activating cAMP signaling has on Mtb metabolism. In this work, we determined that V-59 is an Rv1625c agonist, and its ability to inhibit Mtb growth in macrophages and cholesterol is dependent on Rv1625c and an associated increase in cAMP synthesis. Additionally, we found that the transmembrane domain of Rv1625c is necessary for the complete metabolism of cholesterol, linking the protein target of V-59 directly to the cholesterol utilization pathway. This finding connects a single AC to the regulation of a downstream metabolic pathway in Mtb for the first time. Using a complementary genetic approach, we developed an inducible system to activate cAMP synthesis independent of V-59 and Rv1625c, and determined that upregulating cAMP synthesis is sufficient to inhibit cholesterol utilization in Mtb. V-59 was optimized through medicinal chemistry, which produced a lead compound (mCLB073) with improved potency and *in vivo* activity against Mtb when delivered orally to infected mice. Collectively, our results reveal a novel cAMP signaling mechanism in Mtb that inhibits cholesterol utilization and may represent an improvement over developing conventional single-step inhibitors against this complex pathway. Using a small molecule AC agonist as an antimicrobial compound is an unconventional approach, and this study is the first to explore this as a mechanism of action to inhibit growth of a bacterial pathogen during infection.

## Results

### V-59 inhibits Mtb growth and requires Rv1625c for activity

We previously identified compounds that inhibit Mtb replication in macrophages (*29*) and determined that one of these compounds (V-58) preferentially inhibits Mtb growth in cholesterol media in an Rv1625c dependent manner (*36*). Unfortunately, these previously identified compounds have poor potency and pharmacological properties. For example, a resynthesized analog of V-58 (sCEB942) displayed sub-optimal intramacrophage potency that could not be improved (Supplementary Fig. 1a). Similarly, the previously identified compound (mCCY224) displayed poor solubility, high plasma protein binding, and high levels of caseum binding (Supplementary Fig. 1a). These properties precluded the use of these Rv1625c-dependent compounds in mice. Since a primary goal of this work was to investigate the impact that activating cAMP synthesis has on Mtb physiology during infection in mice, we re-examined our screening hits to identify candidate Rv1625c-dependent compounds with more favorable pharmacological properties that are permissible for both *in vivo* and *in vitro* studies. This effort revealed a small molecule 1-(4-(5-(4-fluorophenyl)-2H-tetrazol-2-yl)piperidin-1-yl)-2-(4-methyl-1,2,5-oxadiazol-3-yl)ethan-1-one, named V-59, that inhibits Mtb replication in macrophages (half maximal effective concentration (EC_50_) 0.30 μM). Because the availability of carbon sources can potentially impact activity of chemical inhibitors against Mtb, V-59 was evaluated in different *in vitro* culture conditions. Similar to a previously characterized Rv1625c agonist (*36*), V-59 inhibits Mtb growth in cholesterol media (EC_50_ 0.70 μM) (Fig. 1a and Table 1) but not in media containing the two-carbon fatty acid acetate, or in standard rich growth media (Fig. 1b and Table 1). V-59 also displayed a promising pharmacokinetic profile (Table 1) and was therefore selected for further investigation as a potential Rv1625c agonist.

**Figure 1.**
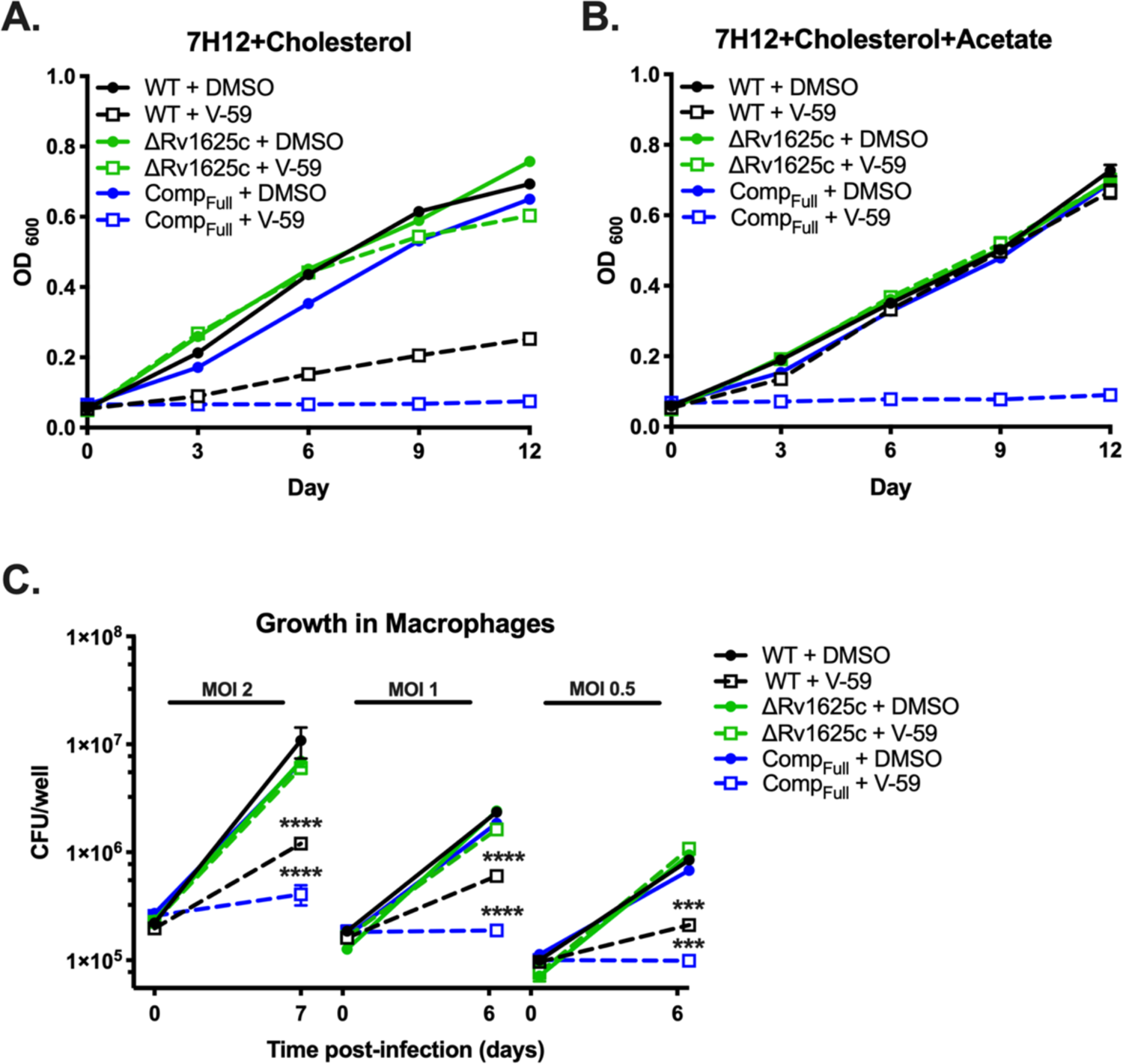
V-59 inhibits Mtb growth in an Rv1625c-dependent mechanism. (**A and B**) Impact of V-59 on Mtb growth in cholesterol media (A) and in media containing cholesterol and acetate (B). V-59 (10 μM) was added to the cultures every three days, and DMSO is the vehicle control. Data are from one experiment, with three technical replicates. (**C**) Effect of V-59 on growth of Mtb in murine macrophages. Macrophages were treated with V-59 (25 μM) or DMSO. Data are from one experiment (MOI 1 and 0.5) with at least two technical replicates, or two experiments (MOI 2) with two technical replicates each (\*\*\**P* < 0.001, \*\*\*\**P* < 0.0001, One-way ANOVA with Sidak’s multiple comparisons test on calculated fold-change in CFU’s). All data are means ± SEM.

**Table 1.**
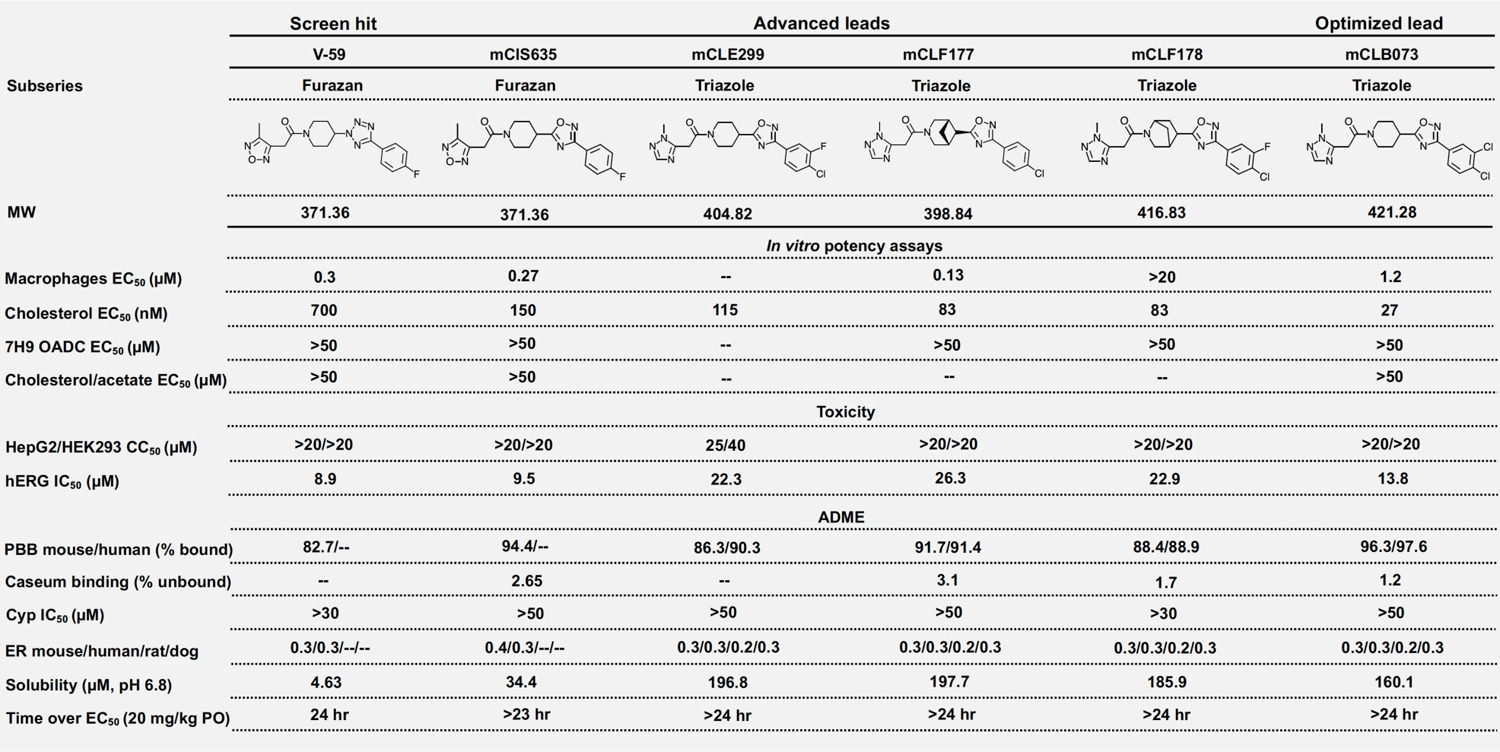
Structures and activities of compounds. MW, molecular weight; --, not determined; EC_50_, half-maximal effective concentration; CC_50_, 50% cytotoxic concentration; hERG, human ether-à-go-go-related gene; IC_50_, half-maximal inhibitory concentration; ADME, absorption, distribution, metabolism, excretion; PPB, plasma protein binding; Cyp, cytochrome P450; ER, extraction ratio; PO, per oral.

| Subseries | Screen hit | Advanced leads |  |  |  | Optimized lead |
| --- | --- | --- | --- | --- | --- | --- |
|  | V-59 | mCIS635 | mCLE299 | mCLF177 | mCLF178 | mCLB073 |
|  | Furazan | Furazan | Triazole | Triazole | Triazole | Triazole |
| MW | 371.36 | 371.36 | 404.82 | 398.84 | 416.83 | 421.28 |
| In vitro potency assays |  |  |  |  |  |  |
| Macrophages EC<sub>50</sub> (μM) | 0.3 | 0.27 | -- | 0.13 | >20 | 1.2 |
| Cholesterol EC<sub>50</sub> (nM) | 700 | 150 | 115 | 83 | 83 | 27 |
| 7H9 OADC EC<sub>50</sub> (μM) | >50 | >50 | -- | >50 | >50 | >50 |
| Cholesterol/acetate EC<sub>50</sub> (μM) | >50 | >50 | -- | -- | -- | >50 |
| Toxicity |  |  |  |  |  |  |
| HepG2/HEK293 CC<sub>50</sub> (μM) | >20/>20 | >20/>20 | 25/40 | >20/>20 | >20/>20 | >20/>20 |
| hERG IC<sub>50</sub> (μM) | 8.9 | 9.5 | 22.3 | 26.3 | 22.9 | 13.8 |
| ADME |  |  |  |  |  |  |
| PBB mouse/human (% bound) | 82.7/-- | 94.4/-- | 86.3/90.3 | 91.7/91.4 | 88.4/88.9 | 96.3/97.6 |
| Caseum binding (% unbound) | -- | 2.65 | -- | 3.1 | 1.7 | 1.2 |
| Cyp IC<sub>50</sub> (μM) | >30 | >50 | >50 | >50 | >30 | >50 |
| ER mouse/human/rat/dog | 0.3/0.3/--/-- | 0.4/0.3/--/-- | 0.3/0.3/0.2/0.3 | 0.3/0.3/0.2/0.3 | 0.3/0.3/0.2/0.3 | 0.3/0.3/0.2/0.3 |
| Solubility (μM, pH 6.8) | 4.63 | 34.4 | 196.8 | 197.7 | 185.9 | 160.1 |
| Time over EC<sub>50</sub> (20 mg/kg PO) | 24 hr | >23 hr | >24 hr | >24 hr | >24 hr | >24 hr |

V-59 is structurally distinct from previous cholesterol utilization inhibitor candidates (Supplementary Fig. 1a). Similar to a subset of other cholesterol-dependent Mtb growth inhibitors we identified (*29*), a transposon insertion in the *rv1625c* gene (Tn::*rv1625c*) confers resistance to V-59 (Supplementary Fig. 1b). Inversely, WT Mtb transformed with an *rv1625c* overexpression plasmid (2x*rv1625c*) was ∼15-fold more susceptible to V-59 than WT (Supplementary Fig. 1b). This heightened susceptibility suggests a mechanism in which V-59 activates Rv1625c, and growth inhibition scales with Rv1625c enzyme levels. To test this further, we deleted the gene encoding Rv1625c (ΔRv1625c) and complemented this mutation with the entire *rv1625c* gene (Comp_Full_). The ΔRv1625c mutant is refractory to V-59 inhibition in cholesterol media (Fig. 1a, Supplementary Fig. 1c). Because macrophages contain various nutrients that can support Mtb growth (*37*) we determined that V-59 inhibits Mtb growth in murine macrophages *in vitro* and confirmed that Rv1625c is required for V-59 activity during macrophage infection (Fig. 1c). Importantly, the ΔRv1625c strain does not have a pan-drug resistance profile (Supplementary Fig. 1d). Across all of these assays, the Comp_Full_ strain was more susceptible to V-59 treatment relative to WT, even in media containing acetate. This is likely because *rv1625c* is overexpressed in the Comp_Full_ strain relative to its native expression levels in WT (Supplementary Fig. 1e). We conclude that a functional Rv1625c enzyme is required for V-59 activity, and that this compound inhibits Mtb growth in cholesterol media and macrophages.

### Rv1625c is necessary and sufficient for V-59 to stimulate cAMP production

Rv1625c is a biochemically confirmed AC enzyme that catalyzes the intramolecular cyclization of ATP into cAMP (*30*). Therefore, we determined whether V-59 increases cAMP production in whole bacteria in an Rv1625c-dependent manner. V-59 induced cAMP by ∼70-fold in WT and ∼140-fold in Comp_Full_, but did not affect the ΔRv1625c mutant (Fig. 2a). To determine whether Rv1625c is sufficient for V-59 to stimulate cAMP production, we heterologously expressed the *rv1625c* gene in an AC-deficient strain of *E. coli*. This strain is deficient in its own single AC (*cya^-^ E. coli*), ensuring that the cAMP produced in this experiment is due to Rv1625c activity (*36*). V-59 treatment significantly increased cAMP levels in *cya*^-^ *E. coli* transformed with the Rv1625c expression plasmid (Fig. 2b). In Mtb, we found that spontaneous mutations in *rv1625c* confer resistance to V-59 (Fig. 2c). Mutations predicted to truncate the Rv1625c protein and inactivate its cyclase domain resulted in resistance (Supplementary Fig. 2a). We also identified missense mutations within the transmembrane and cyclase domains of Rv1625c that confer resistance; without further biochemical characterization, it is ambiguous whether these mutations generate resistance by preventing V-59 binding to Rv1625c, or by disabling Rv1625c enzyme activity. Together these results indicate that V-59 activates Rv1625c selectively in Mtb, and that Rv1625c expression is sufficient for V-59 to activate cAMP synthesis, which is necessary for V-59 to inhibit Mtb growth.

**Figure 2.**
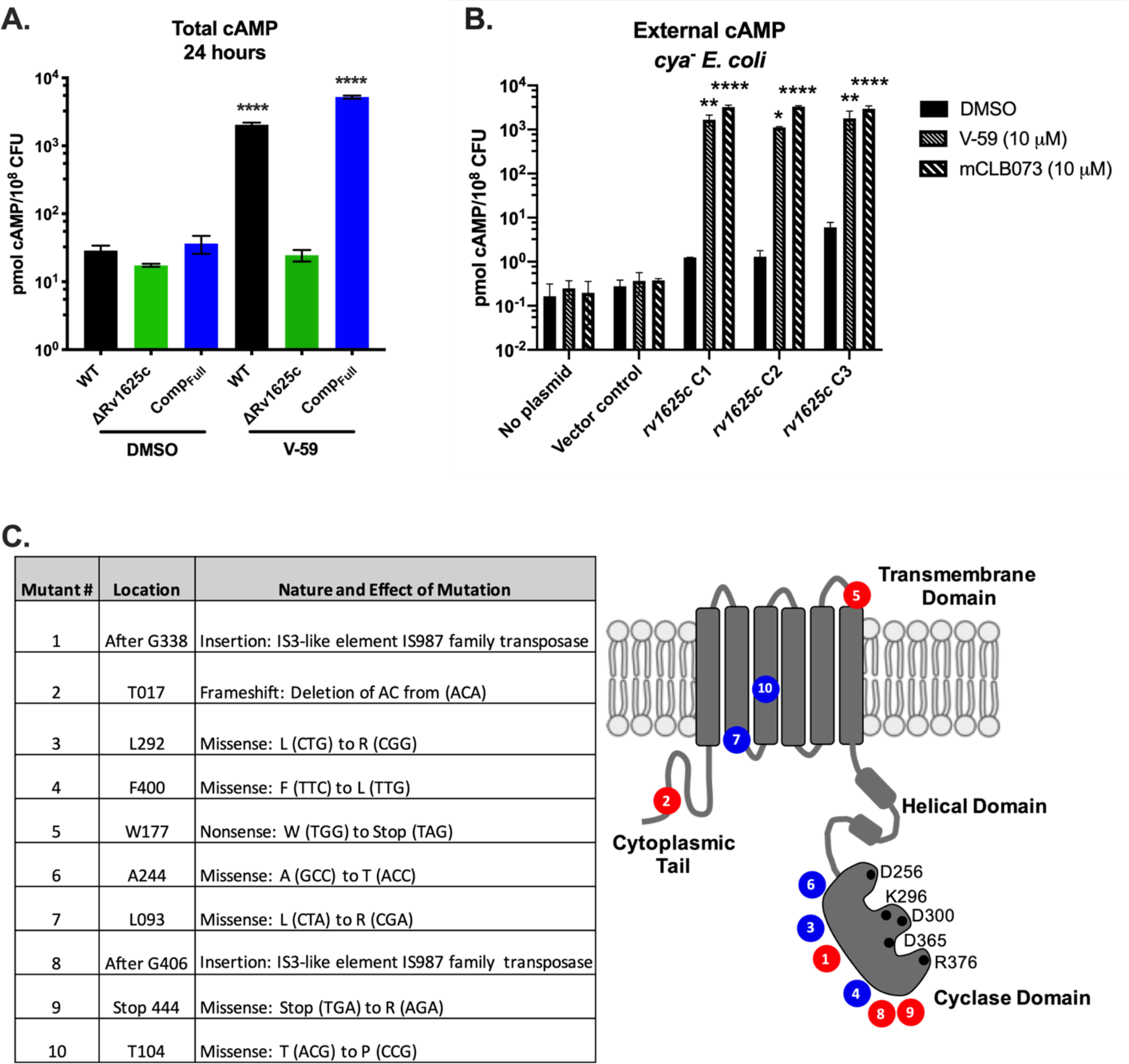
V-59 binds Rv1625c and stimulates cAMP production. (**A**) Impact of V-59 on cAMP production in Mtb. Cultures were treated with V-59 or DMSO for 24 hours. Data are from two experiments with two technical replicates each (\*\*\*\**P* < 0.0001, Two-way ANOVA with Sidak’s multiple comparisons test). (**B**) Impact of Rv1625c agonists on cAMP production in *cya^-^ E. coli* transformed with an empty vector control or an Rv1625c expression plasmid. Supernatants were collected 18 hours after addition of V-59, mCLB073, or DMSO. Data is from one experiment, with three independent expression clones, and two technical replicates each. (\**P* < 0.05, \*\**P* < 0.01, \*\*\*\**P* < 0.0001, Two-way ANOVA with Tukey’s multiple comparisons test). In (A) and (B) Data are normalized as total cAMP per 10^8^ bacteria. DMSO is the vehicle control. Data are shown as means ± SEM. (**C**) Summary of mutations in the *rv1625c* gene that confer resistance to Rv1625c agonists. Mutations are grouped by their effect on the *rv1625c* sequence, with missense mutations (blue) and insertion or frameshift mutations (red) and mapped on the Rv1625c topology diagram to illustrate their approximate location relative to Rv1625c protein domains. Black circles represent amino acids that are essential for AC activity.

### Rv1625c is directly linked to cholesterol degradation in Mtb

Because V-59 impairs growth of Mtb in cholesterol media, we tested whether using V-59 to chemically activate Rv1625c inhibits the bacterium’s ability to break down cholesterol. When Mtb degrades the A-ring of [4-^14^C]-cholesterol, [1-^14^C]-pyruvate is released; subsequently, pyruvate dehydrogenase activity mediates the conversion of [1-^14^C]-pyruvate into acetyl-CoA and ^14^CO_2_ (*6*). Therefore, to quantify cholesterol degradation in Mtb, we captured ^14^CO_2_ released following [4-^14^C]-cholesterol breakdown by the bacteria (*38*). We found that V-59 decreased ^14^CO_2_ release in WT by ∼89% (Fig. 3a). By contrast, V-59 had no measurable effect on ^14^CO_2_ released from breakdown of the fatty acid [U-^14^C]-palmitate (Supplementary Fig. 3a). This suggests that chemically activating Rv1625c preferentially inhibits cholesterol utilization in WT Mtb, rather than equally inhibiting all lipid utilization by the bacterium.

**Figure 3.**
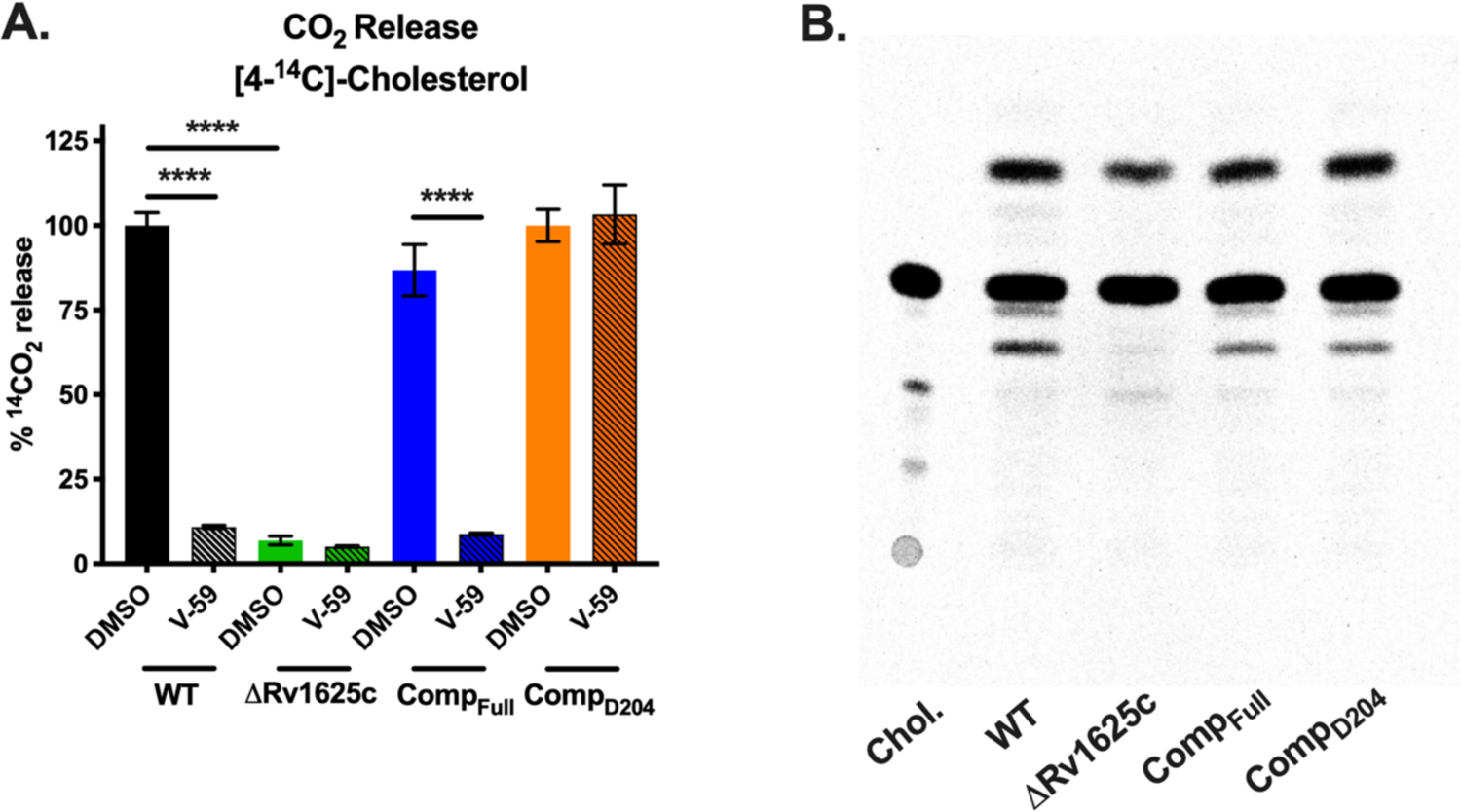
The transmembrane domain of Rv1625c is essential for complete degradation of cholesterol and the catalytic domain of Rv1625c is required for V-59 activity. (**A**) Catabolic release of ^14^CO_2_ from [4-^14^C]-cholesterol in WT, ΔRv1625c, Comp_Full_, and Comp_D204_ strains treated with V-59 (10 μM) or DMSO vehicle control. Data are from two experiments with three technical replicates, normalized to OD and quantified relative to WT treated with DMSO. Shown as means ± SEM (\*\*\*\**P* < 0.0001, Two-way ANOVA with Tukey’s multiple comparisons test). (**B**) TLC comparing [4-^14^C]-cholesterol-derived metabolites extracted from supernatants of Mtb. Image is representative of two experiments. Equivalent counts were spotted per lane. “Chol.” = [4-^14^C]-cholesterol.

Unexpectedly, we found that the ΔRv1625c mutant has an intrinsic defect in cholesterol degradation (Fig. 3a). In contrast to ΔRv1625c, the Rv1625c transposon mutant strain (Tn::*rv1625c*) had no defect in ^14^CO_2_ release from [4-^14^C]-cholesterol (Supplementary Fig. 3b). The Tn::*rv1625c* strain has a transposon insertion within the coding sequence located after the last exit of Rv1625’s six-helical transmembrane domain (amino acid Y302) (Supplementary Fig. 3c). This likely truncates the protein, eliminating more than half of the C-terminal cyclase domain, while leaving the N-terminal cytoplasmic tail and six-helical transmembrane domain intact. Thus, we complemented the ΔRv1625c strain with a construct that expresses only the N-terminal cytoplasmic tail and six-helical transmembrane domain of Rv1625c (Comp_D204_) (Supplementary Fig. 3c). Cholesterol degradation was restored in the Comp_D204_ strain (Fig. 3a), indicating that the transmembrane domain of Rv1625c is required for the complete degradation of cholesterol.

Importantly, V-59 inhibited ^14^CO_2_ release in the Comp_Full_ strain; however, V-59 did not prevent ^14^CO_2_ release in the Comp_D204_ strain which lacks the Rv1625c cyclase domain (Fig. 3a).

To further examine whether cholesterol degradation is blocked in ΔRv1625c Mtb, we used thin-layer chromatography (TLC) to track accumulation of [4-^14^C]-cholesterol-derived metabolites. Compared to WT Mtb, the culture supernatant of ΔRv1625c was deficient in at least one cholesterol-derived degradation intermediate, and the production of this intermediate was restored in the Comp_Full_ strain (Fig. 3b). Collectively, these results indicate that the cyclase domain of Rv1625c must be present for V-59 to inhibit cholesterol catabolism, and the transmembrane domain of Rv1625c is required for complete cholesterol breakdown, thereby establishing a direct link between the target of V-59 and the cholesterol pathway in Mtb. To our knowledge, this is the first time an individual AC has been linked to modulation of a downstream metabolic pathway in Mtb.

### Inducing cAMP synthesis is sufficient to regulate cholesterol utilization

Next, we investigated whether cAMP signaling can modulate cholesterol metabolism in an Rv1625c-independent manner by using a novel inducible construct (TetOn-cAMP) to increase cAMP synthesis in Mtb. This TetOn-cAMP construct carries an anhydrotetracycline (Atc) inducible promoter that controls expression of the catalytic domain of the mycobacterial AC Rv1264 (*18*) (Supplementary Fig. 4a). Atc induced cAMP synthesis in WT Mtb carrying the TetOn-cAMP construct in a dose-dependent manner, reaching levels comparable with V-59 treatment (Fig. 4a). This tool is an advancement over previous approaches (*24, 39*) for several reasons: it does not rely on diffusion of an external cAMP analog into the bacteria, it requires the bacteria to synthesize cAMP from ATP which more closely models the dynamics of AC signaling, and it increases cAMP by 24 hours post-induction in a dose-dependent fashion. Atc treatment inhibited growth of WT bacteria carrying the TetOn-cAMP construct in cholesterol media (Fig 4b) and also decreased [4-^14^C]-cholesterol degradation to ^14^CO_2_ (Fig. 4c). Similar to V-59 treatment, activating cAMP synthesis with Atc did not inhibit degradation of the fatty acid [U-^14^C]-palmitate to ^14^CO_2_ (Supplementary Fig. 3a). As a control, we modified the TetOn-cAMP construct by mutating a catalytic residue of Rv1264 (TetOn-Rv1264_D265A_) to render it catalytically inactive (*18*). Atc induced expression of the Rv1264_D265A_ protein in WT Mtb carrying the TetOn-Rv1264_D265A_ construct (Supplementary Fig. 4b), but this strain did not produce increased cAMP in response to Atc (Supplementary Fig. 4c). Inducing Rv1264_D265A_ expression did not inhibit bacterial growth in cholesterol media (Supplementary Fig. 4d) or cholesterol degradation (Supplementary Fig. 4e, f). These results demonstrate that activating cAMP synthesis through a mechanism that is independent of Rv1625c is sufficient to regulate cholesterol utilization in Mtb in a dose-dependent manner.

**Figure 4.**
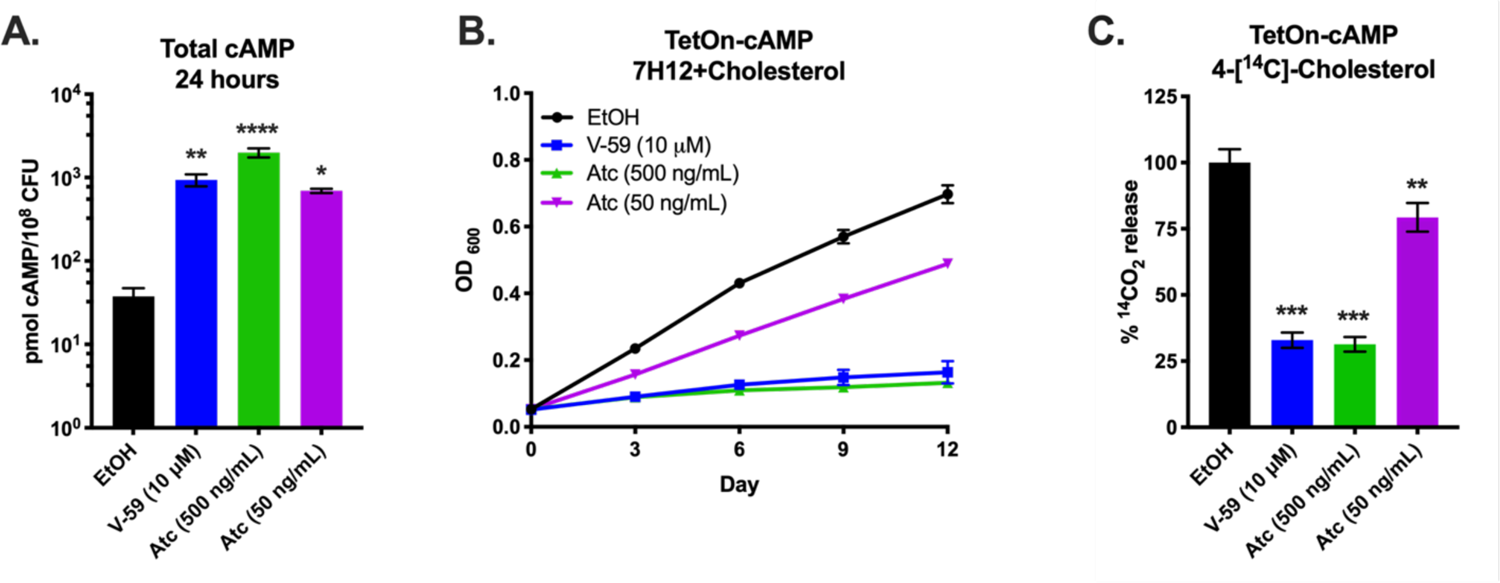
Inducing cAMP synthesis independent of V-59 and Rv1625c is sufficient to block cholesterol utilization. (**A**) Total cAMP induced in TetOn-cAMP Mtb. Cultures were treated with V-59 (10 μΜ), Atc (500 ng/mL or 50 ng/mL), or EtOH and samples were collected after 24 hours. Data are normalized as total cAMP per 10^8^ Mtb and are from two experiments with two technical replicates each (\**P* < 0.05, \*\**P* < 0.01, \*\*\*\**P* < 0.0001, One-way ANOVA with Dunnet’s multiple comparisons test). (**B**) Impact of inducing TetOn-cAMP on the growth of Mtb in cholesterol media. Cultures were treated with V-59 (10 μM) or Atc for the duration of the experiment. Data are from two experiments with three technical replicates. (**C**) Catabolic release of ^14^CO_2_ from [4-^14^C]-cholesterol in the TetOn-cAMP strain treated with V-59, Atc, or EtOH. Data are from two experiments with three technical replicates, normalized to OD and quantified relative to EtOH (\*\**P* < 0.01, \*\*\**P* < 0.001, One-way ANOVA with Dunnet’s multiple comparisons test). EtOH is the vehicle control throughout. All data are means ± SEM.

### Common transcriptional changes in cholesterol genes are associated with cAMP induction

Next, we characterized transcriptional responses of Mtb following V-59 treatment, or upon induction of the TetOn-cAMP construct, during growth in cholesterol media. The RNA-seq datasets revealed a shared pattern of differential gene expression that is consistent with an early blockade in the cholesterol degradation pathway. In Mtb, the side-chain and A-B rings of cholesterol are degraded by enzymes encoded in the Rv3574/KstR1 regulon (*6*). KstR1 is a TetR-like transcriptional repressor that binds the second cholesterol degradation intermediate, 3-hydroxy-cholest-5-ene-26-oyl-CoA, which de-represses the KstR1 regulon and permits cholesterol degradation to occur. Thus, increased expression of the KstR1 regulon is an indicator of cholesterol degradation in Mtb. Inducing cAMP synthesis, with V-59 or by activating TetOn-cAMP, prevented transcriptional induction of the KstR1 regulon in WT Mtb (Fig. 5). This included key genes required for cholesterol transport (*rv0655/mceG* and *rv3502/yrbE4B*) and cholesterol catabolism (*6*). Cholesterol degradation releases propionyl-CoA, and Mtb primarily assimilates this intermediate into central metabolism via the methylcitrate cycle (MCC) (*6*). As propionyl-CoA pools increase, Mtb upregulates expression of the genes encoding MCC enzymes (*rv0467/icl1*, *rv1130/prpD*, *rv1131/prpC*) (*40*). V-59 treatment and induction of the TetOn-cAMP construct in WT Mtb each prevented upregulation of MCC genes (Fig. 5). Paralleling previous experiments, V-59 induced more pronounced changes in the transcriptional signature in the Comp_Full_ strain. Notably, genes required for cholesterol transport and genes (*hsaEFG*) necessary for conversion of the cholesterol-derived catabolic intermediate 2-hydroxy-hexa-2,4-dienoic acid to pyruvate and propionyl-CoA were upregulated in the Comp_Full_ strain following V-59 treatment (*6*). It is plausible that these expression profiles reflect a compensatory response to inhibition of cholesterol degradation by V-59, and a concomitant decrease in availability of MCC or tricarboxylic acid cycle intermediates. Consistent with our previous observations (Fig. 3), expression of cholesterol side-chain and ring degradation genes, but not transport or MCC genes, was intrinsically blocked in the ΔRv1625c strain relative to WT (Fig. 5). These observations further support the conclusion that Rv1625c is involved in downstream cholesterol metabolism in Mtb. Importantly, V-59 treatment did not alter the transcriptional signature of the ΔRv1625c strain.

**Figure 5.**
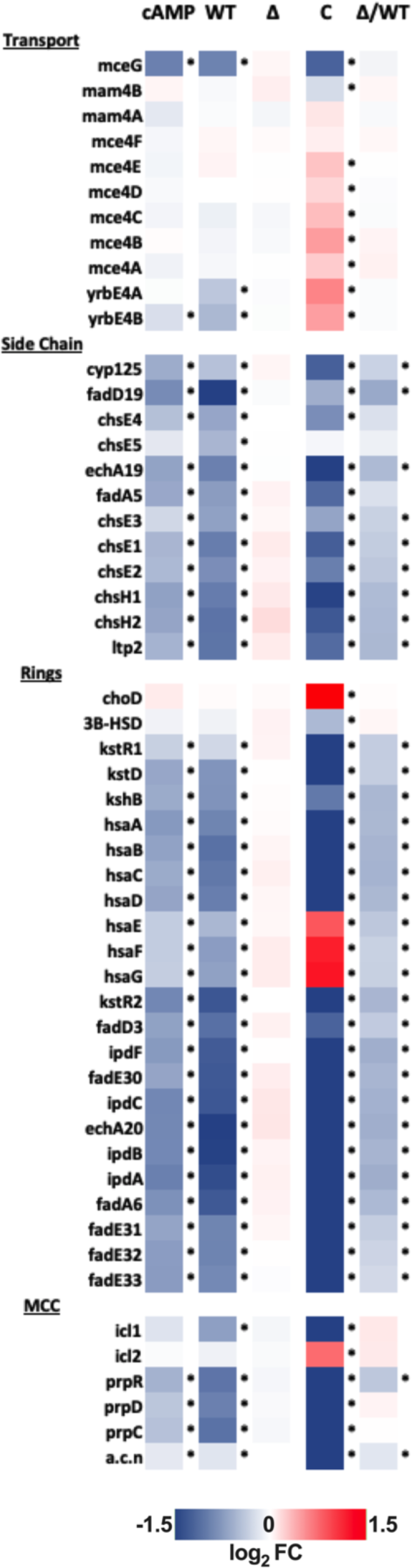
V-59 treatment and induction of TetOn-cAMP are associated with shared transcriptional changes to cholesterol utilization genes. RNA-seq analysis quantifying differentially expressed genes from Mtb grown in cholesterol media, following V-59 treatment, or induction of TetOn-cAMP with Atc. Genes depicted are in the KstR regulons and involved in cholesterol utilization. MCC = methylcitrate cycle. Data are displayed as log_2_ fold change in gene expression in response to cAMP-inducing vs. control treatment (“cAMP” = Tet-On cAMP Atc vs. EtOH, “WT” = WT V-59 vs. DMSO, “Δ” = ΔRv1625 V-59 vs. DMSO, “C” = Comp_Full_ V-59 vs. DMSO). Also shown are differentially expressed genes intrinsic to ΔRv1625 (“Δ/WT” = ΔRv1625 DMSO vs. WT DMSO). Data are from two technical replicate samples from one experiment (*adjusted p-value ≤ 0.05).

To validate these findings, we used a reporter (*prpD’*::GFP) that expresses GFP under control of a MCC gene promoter (*prpD*), which indicates cellular levels of propionyl-CoA (*38*). V-59 decreased GFP signal in WT by ∼50%, but did not impact the ΔRv1625c or Comp_D204_ strains in cholesterol media or during macrophage infection (Fig. 6a). V-59 also dampened GFP signal by ∼90% in the Comp_Full_ strain (Fig. 6a). Similarly, inducing cAMP synthesis in WT Mtb carrying the TetOn-cAMP construct was sufficient to inhibit GFP signal during growth in cholesterol media and during macrophage infection (Fig. 6b). Inducing expression of the inactive Rv1264_D265A_ protein (Supplementary Fig. 4b) did not change the GFP signal (Supplementary Fig. 4f). These data demonstrate that inducing cAMP synthesis in Mtb, via V-59 treatment or TetOn-cAMP activation, impairs cholesterol degradation and the release of key metabolic intermediates including propionyl-CoA in Mtb. Importantly, the effects of V-59 treatment require the catalytic domain of Rv1625c, and cAMP synthesis is a dominant signal in the mechanism by which V-59 inhibits Mtb growth.

**Figure 6.**
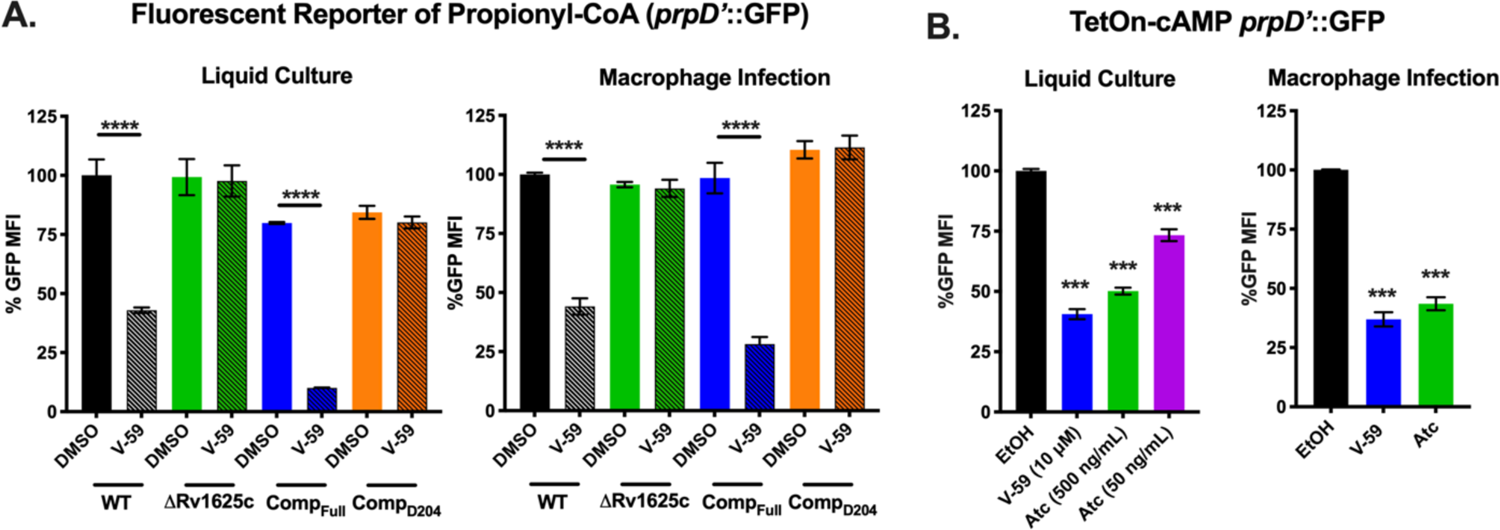
Activating cAMP synthesis decreases liberation of propionyl-CoA from cholesterol. (**A**) Relative GFP signal from the *prpD’*::GFP reporter in response to V-59 (10 μM) or DMSO treatment in murine macrophages or cholesterol media. Data are normalized to WT treated with DMSO (\*\*\*\**P* < 0.0001, Two-way ANOVA with Tukey’s multiple comparisons test). (**B**) Relative GFP signal from the *prpD’*::GFP reporter in response to inducing TetOn-cAMP with Atc treatment in murine macrophages or cholesterol media. Data are normalized to EtOH vehicle control (\*\*\**P* < 0.001, One-way ANOVA with Dunnett’s multiple comparisons test). GFP MFI was quantified from 10,000 mCherry^+^ Mtb. Data are from two experiments with two technical replicates, shown as means ± SEM.

### Mt-Pat is not required to mediate inhibition of cholesterol utilization

Inducing cAMP synthesis blocks cholesterol utilization in Mtb, but the mechanism mediating this is unknown. Because fatty acid metabolism can be modulated by the cAMP-binding protein Rv0998/Mt-Pat (*27, 28*) we investigated whether Mt-Pat also mediates V-59-dependent inhibition of cholesterol utilization. However, inhibition of growth (Supplementary Fig. 5a) and inhibition of MCC gene induction (Supplementary Fig. 5b) by V-59 treatment were not altered in an Mt-Pat mutant. Notably, the Comp_Full_ strain was uniquely susceptible to V-59 in cholesterol media supplemented with the short chain fatty acid acetate (Fig. 1b, Supplementary Fig. 5c), and V-59 was also found to block MCC gene induction in the Comp_Full_ strain during growth with odd-chain fatty acids (Supplementary Fig. 5d). This suggests that additional metabolic defects, possibly in fatty acid utilization or central metabolism, are induced under these conditions. While these observations correlate with a higher threshold of cAMP induction (Fig. 2a), we have not determined whether Mt-Pat mediates these additional effects. In the future, identifying the pathway by which inducing cAMP synthesis modulates cholesterol catabolism in Mtb may explain the differing effects of V-59 on carbon metabolism in these strains.

### Transcriptional changes in select CRP_MT_ regulon genes are associated with cAMP induction

To define a set of commonly regulated cAMP-dependent genes using an unbiased analysis, we compared all of the statistically significant differentially expressed genes associated with V-59 treatment or TetOn-cAMP induction in WT Mtb (Fig. 7a). A shared set of 248 genes were identified (Fig. 7b). Because the selected growth condition was cholesterol media, 45 of these genes are associated with cholesterol utilization. As discussed above, those with biochemically confirmed roles in cholesterol utilization were noted (Fig. 5). Many of the remaining 203 genes do not have well defined functions in Mtb, and we chose to focus on a small subset that were previously predicted to be regulated by the cAMP-binding transcription factor Rv3676/CRP_MT_.

**Figure 7.**
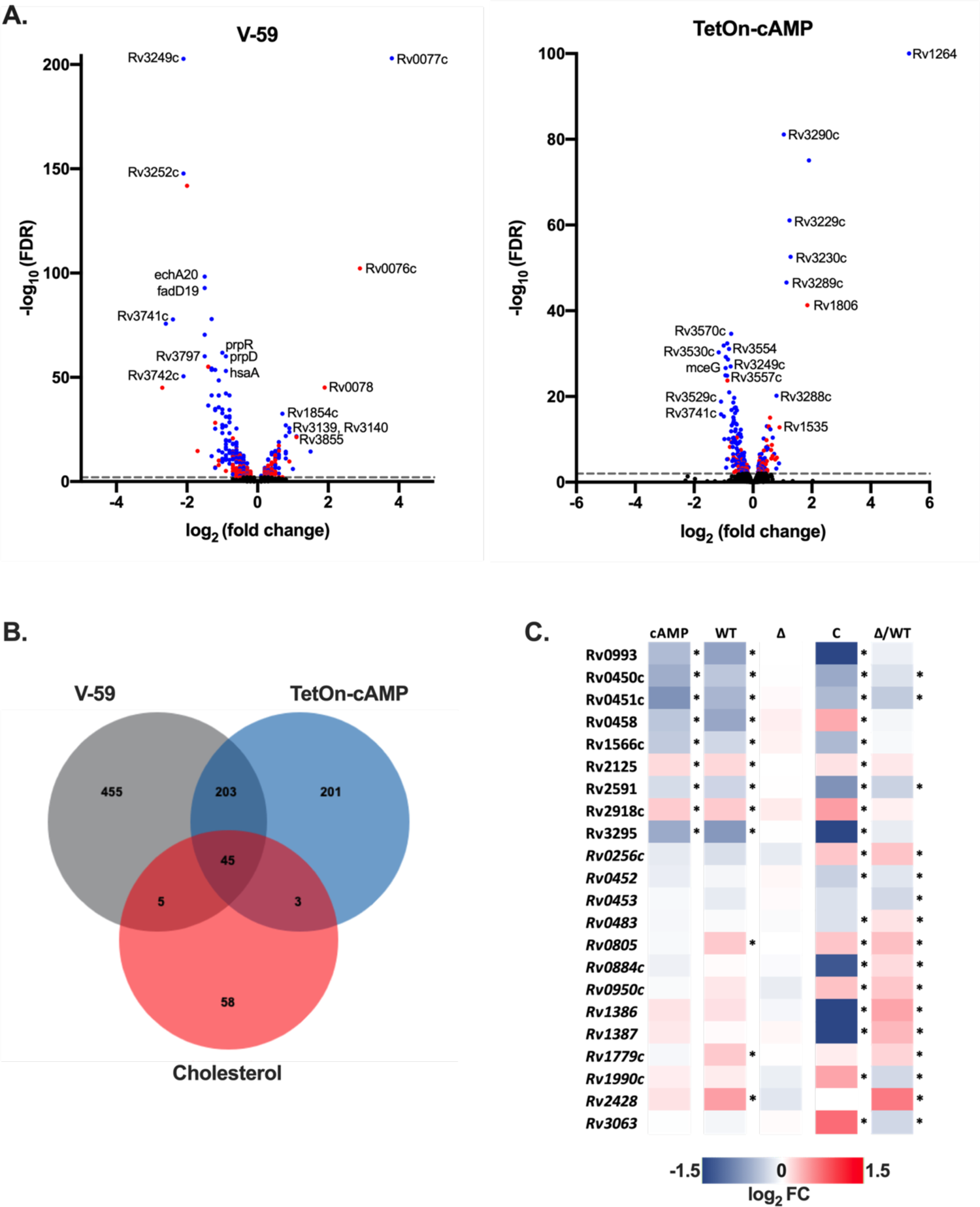
V-59 treatment and induction of TetOn-cAMP are associated with transcriptional changes in select CRP_Mt_ regulon genes. (**A**) Volcano plots displaying differentially expressed genes following V-59 treatment of WT Mtb relative to DMSO control (left), or following Atc treatment of TetOn-cAMP Mtb relative to EtOH control (right), based on RNA-seq. Each dot represents a single gene, genes in blue are significant (FDR < 0.05) in both data sets and genes in red are unique to their respective data set. Dashed line indicates FDR cutoff < 0.01. (**B**) Venn diagram showing the number of significantly differentially expressed genes shared by the V-59 and TetOn-cAMP conditions, and how many of these belong to the KstR cholesterol-related regulon. (**C**) RNA-seq analysis quantifying differentially expressed genes from Mtb grown in cholesterol media, following V-59 treatment, or induction of TetOn-cAMP with Atc. Genes depicted are predicted members of the CRP_MT_ regulon. Only genes with significant differential expression in both the V-59 and TetOn-cAMP conditions, or genes with intrinsic changes in the ΔRv1625 strain (in italics), are shown. “cAMP” = Tet-On cAMP Atc vs. EtOH, “WT” = WT V-59 vs. DMSO, “Δ” = ΔRv1625 V-59 vs. DMSO, “C” = Comp_Full_ V-59 vs. DMSO, “Δ/WT” = ΔRv1625 DMSO vs. WT DMSO.

Aside from Mt-Pat, CRP_Mt_ is the best-studied cAMP-binding effector protein in Mtb. CRP_Mt_ is designated as a cAMP-responsive transcription factor, with a predicted regulon of ∼100 genes in Mtb (*25, 26, 41*). CRP_Mt_ may also be required to maintain Mtb fitness in macrophages and during chronic infection in mice (*26*). We found that activating cAMP via V-59 treatment or TetOn-cAMP induction was associated with transcriptional changes to a shared set of 8 CRP_Mt_ regulon genes during growth in cholesterol media (Fig. 7c). Among these, *rv0450c/mmpL4* and *rv0451c/mmpS4* are both downregulated following cAMP induction. Though *mmpL4* was not a predicted member of the CRP_MT_ regulon, it is reasonable that these genes would be co-regulated given that MmpL4 and MmpS4 are encoded in the same putative operon and can form a protein complex that contributes to siderophore production in Mtb (*42*). This result is also notable because genetic screens have predicted that *mmpL4/mmpS4* are required for normal growth of Mtb in cholesterol media and in mouse models of TB (*12, 43*). We also noted that the predicted CRP_MT_ regulon member *rv0805* is upregulated during V-59 treatment. Rv0805 is the only known phosphodiesterase in Mtb, and these enzymes contribute to cAMP signaling pathway homeostasis by hydrolyzing cAMP to AMP (*23*). Given that *rv0805* mutant Mtb also has a growth defect in cholesterol media (*43*), it is reasonable to speculate that *rv0805* is upregulated during V-59 treatment in the presence of cholesterol as a compensatory response to help decrease cAMP levels and restore cholesterol utilization. Taken together, this is consistent with our other results indicating that a threshold of increased cAMP is inhibitory during cholesterol utilization in Mtb.

Surprisingly, an additional set of 13 predicted CRP_Mt_ regulon genes displayed intrinsic differential expression in the ΔRv1625c strain relative to WT (Fig. 7c) but were not universally differentially expressed in response to cAMP induction. This suggests that Rv1625c might play a native role in regulating some CRP_MT_ operon genes during cholesterol utilization. Overall, these findings are significant because they demonstrate that a subset of predicted CRP_Mt_ genes are altered either in response to induction of cAMP synthesis, or through loss of Rv1625c, in the presence of cholesterol. While this study does not explain the native role of CRP_MT_ during infection, V-59 and TetOn-cAMP can be used as tools in future studies to examine regulation of this operon under different growth conditions or during infection which may provide insight into its function in Mtb pathogenesis.

### mCLB073 is an optimized analog of the V-59 compound series

Next, we sought to identify chemical features that are essential to a potent Rv1625c agonist, and to develop an optimized compound for use during *in vivo* studies. The screening hit (V-59) was relatively potent against Mtb in macrophages and had several satisfactory pharmacological features including plasma exposure above the EC_50_ (as determined in cholesterol media) for approximately 24 hours following oral dosing in mice at 20 mg/kg (Table 1). We sought to improve the properties of the V-59 compound series with medicinal chemistry. Structure activity relationship studies determined that replacing the tetrazole ring in V-59 with the oxadiazole ring in mCIS635 improved potency and slightly improved solubility. Replacing the 4-methyl-1,2,5-oxadiazole ring in V-59 with a 1-methyl-1H-1,2,4-triazole ring addressed the liability of the oxadiazole ring and generated the lead compounds mCLE299, mCLF177, mCLF178, and mCLB073 that had improved properties including better potency and extended plasma exposure following oral administration in mice. We explored constraining the piperidine ring in order to increase compound solubility by lowering its lattice energy, through an azabicyclic ring and a chiral center in several molecules of this series (Table 1 and Table S1). Interestingly, the *cis* isomer (mCLF024) displayed potency similar to the advanced lead compounds in this series, while the *trans* isomer (mCLF025) was inactive (Table S1).

Among the optimized analogs, the lead compound (1-(4-(3-(3,4-dichlorophenyl)-1,2,4-oxadiazol-5-yl)-1-piperidinyl)-2-(2-methyl-2H-1,2,4-triazol-3-yl)-1-ethanone), named mCLB073, exhibited a ∼17-fold potency improvement against Mtb in cholesterol media relative to V-59 while maintaining excellent pharmacokinetic properties and a good safety profile. We then verified that mCLB073 retained on-target activity. The ΔRv1625c strain was refractory to mCLB073 treatment (Supplementary Fig. 2b), and mCLB073 activates cAMP synthesis (Fig. 2b and Supplementary Fig. 2c). Additionally, we isolated spontaneous resistant mutants in Mtb cultured with mCLB073. All spontaneous resistant mutants we isolated contained mutations in *rv1625c* that conferred resistance to mCLB073 and cross-resistance to V-59 (Fig. 2c and Supplementary Fig. 2a). These results indicate that mCLB073 is a genuine Rv1625c agonist. When dosed orally in mice, mCLB073 maintained plasma exposure over the EC_50_ identified in cholesterol media for at least 24 hours (Table 1). This demonstrated that mCLB073 is suitable for once-daily oral dosing in mouse models of infection and justified using this compound as a chemical probe during *in vivo* studies.

### Selective activation of cAMP synthesis inhibits Mtb pathogenesis in vivo

Next, we examined whether treatment with Rv1625c agonists would alter Mtb survival in a mouse model of infection. We infected BALB/c mice with WT Mtb, and administered a vehicle control, V-59, or isoniazid by oral gavage once-daily during weeks 4 through 8 post-infection. V-59 (50 mg/kg) caused a ∼0.4-log_10_ reduction in lung CFU’s and reduced the extent of lung inflammation by ∼50% (Fig. 8a, b). Similar results were obtained in CFU counts (∼0.5-log_10_ reduction) in the lungs of C3HeB/FeJ mice infected and treated in the same manner (Fig. 8c, d). The C3HeB/FeJ Mtb infection model was used because these mice produce type I IFN- and neutrophil-driven pathology that results in well-organized necrotic granulomas containing high numbers of extracellular bacterial (*44–47*). Thus, the finding that an Rv1625c agonist inhibits bacterial growth and limits lung pathology in both a relatively resistant and a susceptible model of TB suggests that this compound series could be effective despite the heterogeneous host response mounted in Mtb infections (*48*). To verify the improved potency of mCLB073 against Mtb *in vivo*, we infected BALB/c mice with WT Mtb via the intranasal route, and administered a vehicle control, mCLB073, or isoniazid by oral gavage once-daily during weeks 4 through 8 post-infection. Treatment with mCLB073 (30mg/kg) reduced Mtb CFUs in the lungs of mice significantly (∼0.4-log_10_ reduction) (Fig. 8e) and decreased the extent of lung pathology by ∼45% (Fig. 8f). In a separate study of BALB/c mice that were aerosol infected and treated in the same manner, we observed a significant reduction in lung CFUs (∼0.4-log_10_ reduction) at a lower dose of mCLB073 (5mg/kg) which further confirms the improved pharmacological properties of mCLB073 (Fig. 8g).

**Figure 8.**
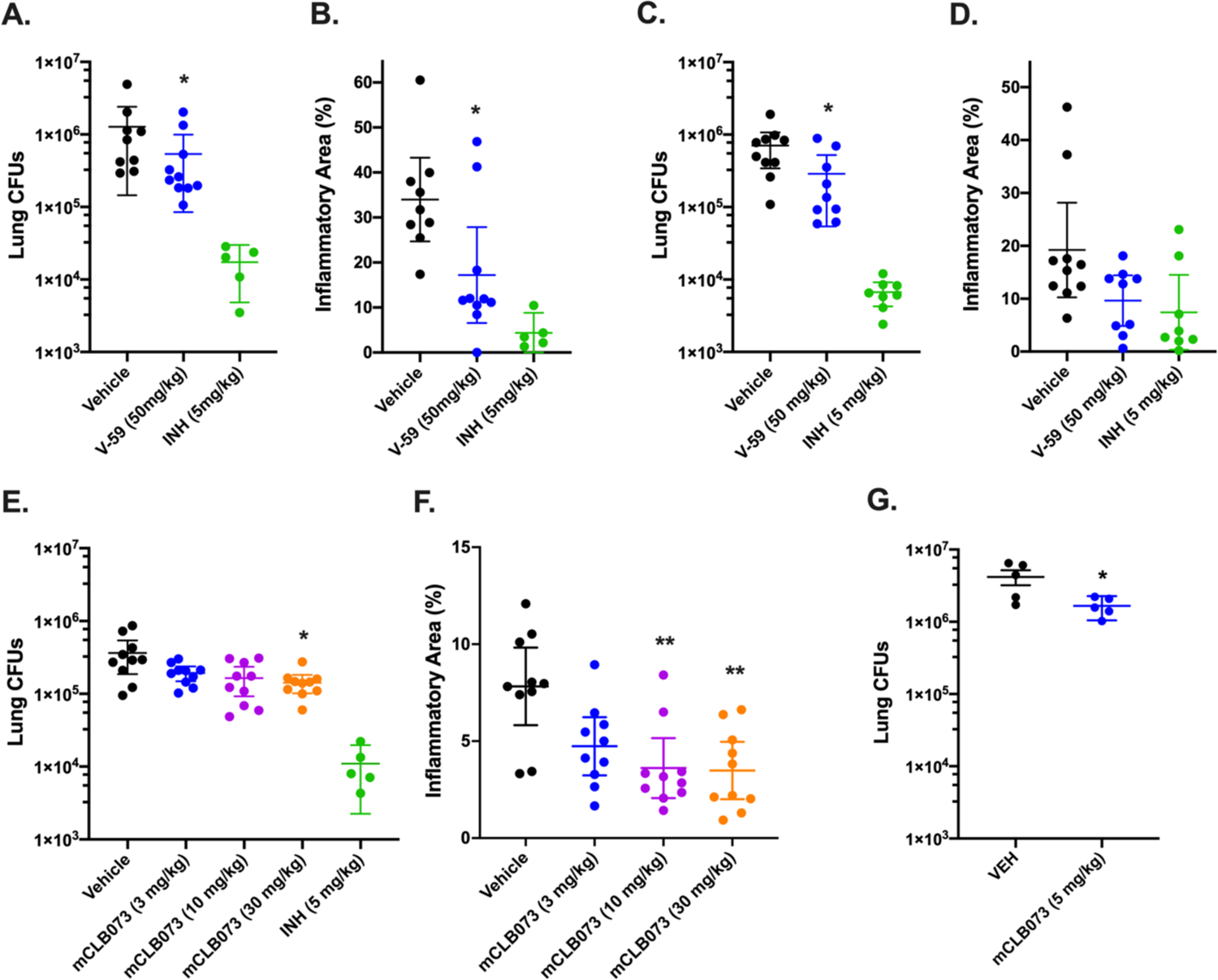
Chemically activating Rv1625c reduces Mtb pathogenesis *in vivo.* Effect of V-59 treatment on bacterial burden and pathology in the lungs of BALB/c (**A and B**) or C3HeB/FeJ mice (**C and D**). In (A-D) mice were infected and treated with V-59, INH, or vehicle control. Data are from two independent experiments with 5 mice (A and B), or one experiment with 10 mice (C and D) per group. Outliers with CFUs below the infectious dose were excluded from the analyses (\**P* < 0.05, Mann-Whitney test). (**E and F**) Impact of mCLB073 treatment on bacterial burden (E) and pathology (F) in BALB/c mice infected and treated with the indicated doses of mCLB073, INH, or vehicle control (\**P* < 0.05, \*\**P* < 0.01, Kruskal-Wallis test and Dunn’s multiple comparisons test). Infections in (A-F) were by the intranasal route. Data are from one experiment with 10 mice per group. (**G**) Impact of mCLB073 treatment on bacterial burden in BALB/c mice infected by aerosol and treated with 5mg/kg mCLB073 or vehicle control. Data are from one experiment with 5 mice per group (\**P* < 0.05, Kruskal-Wallis test and Dunn’s multiple comparisons test). All data are shown as means ± SEM.

Because the mechanism of action of mCLB073 is novel, we tested whether mCLB073 treatment would lead to increased tolerance to a frontline TB drug (rifampicin) during infection. We found that the addition of mCLB073 (30 mg/kg) to a sub-optimal dose of rifampicin did not increase the bacterial burden in the lungs of BALB/c mice, suggesting this mechanism of action does not promote tolerance to other TB antibiotics (Supplementary Fig. 5e). Finally, we addressed the potential concern that this chemotype would activate off-target, mammalian AC enzymes. We found no evidence that V-59 activates ACs in mammalian cells (Supplementary Fig. 5f), and the low toxicity profile of the Rv1625c-activating compounds (Table 1) suggests limited off-target activation of mammalian ACs. These results demonstrate the increased potency of mCLB073 relative to V-59 *in vivo*, and suggest that chemically activating cAMP synthesis in Mtb during chronic infection confers a fitness cost to the bacterium.

## Discussion

Mtb possesses an expanded repertoire of cAMP signaling pathway components compared to other bacteria, suggesting this is an important mechanism to coordinate physiological functions in response to environmental cues. However, how Mtb physiology can be regulated through cAMP signaling, particularly through activation of specific AC enzymes, is not well understood. It was also not previously established whether this signaling pathway could be manipulated pharmacologically to disrupt Mtb pathogenesis. This gap in knowledge is partly explained by the lack of chemical and genetic tools that are equivalent to the eukaryotic AC agonist forskolin in the Mtb system. In this study we identified a chemical AC agonist that is suitable for *in vitro* and *in vivo* studies. And in a complementary approach, we created and validated a TetOn-cAMP construct that permits dose-dependent induction of cAMP synthesis in Mtb. These tools allowed us to establish a link between induction of cAMP synthesis, downregulation of cholesterol utilization, and inhibition of Mtb pathogenesis during infection.

Here, we re-examined a collection of compounds that were identified in a high-throughput screen as inhibitors of Mtb growth in macrophages and cholesterol media. Based on a previous study (*29*) we knew that this collection contained at least three Rv1625c-dependent compounds, and sought to identify an additional Rv1625c agonist with improved potency and acceptable properties for use in *in vivo* studies. From the screening hits, we identified a candidate small molecule named V-59 that displayed promising pharmacological properties (Table 1). We then determined that growth inhibition by V-59 in cholesterol media and in macrophages requires a functional Rv1625c enzyme (Fig. 1a, c and Fig. 2c), and V-59 induces cAMP synthesis in an Rv1625c-dependent manner (Fig. 2a, b). By quantifying degradation of the A-ring of cholesterol, we found that V-59 indeed blocks cholesterol utilization, in a mechanism that requires the cyclase domain of Rv1625c (Fig. 3). Our combined results support the conclusion that V-59 directly binds to the Rv1625c enzyme to activate AC activity, which artificially increases cAMP synthesis in Mtb to inhibit cholesterol utilization which impairs bacterial growth in cholesterol media and macrophages.

To study the impact of cAMP induction independent of V-59 and Rv1625c, we developed a TetOn-cAMP construct that increases cAMP synthesis in a dose-dependent manner (Fig. 4a). We found that inducing cAMP synthesis is sufficient to decrease growth of Mtb in cholesterol media (Fig. 4b) and to block cholesterol degradation (Fig. 4c). Transcriptional studies revealed hallmarks indicating that cholesterol degradation is inhibited early in the breakdown process following V-59 treatment, or induction of cAMP synthesis via the TetOn-cAMP construct (Fig. 5 and Fig. 6). Together, this demonstrates that inducing cAMP above a certain threshold through a different AC is sufficient to mimic the effects of an Rv1625c agonist, which suggests that AC activation is a general mechanism that can be leveraged to inhibit cholesterol utilization in Mtb. Collectively, our findings indicate that inducing cAMP synthesis in Mtb inhibits cholesterol degradation, blocks transcriptional activation of hallmark cholesterol utilization genes, and decreases propionyl-CoA pools in proportion to the amount of cAMP induced. Cholesterol breakdown is a many-stage process, and side chain and ring degradation can occur in tandem (*6*). Considering this, one interpretation consistent across these results is that robust activation of cAMP synthesis prevents cholesterol side chain and ring degradation simultaneously, and this decreases the breakdown of cholesterol to an early intermediate that is required to de-repress the KstR1 regulon.

While investigating the effect of V-59 on Rv1625c activity and cholesterol utilization, we unexpectedly discovered that the six-helical transmembrane domain of Rv1625c and the associated N-terminal cytoplasmic tail is intrinsically required for complete cholesterol degradation in Mtb (Fig. 3a, b). Given that Rv1625c had no previously predicted role in cholesterol utilization, this is a surprising connection between the AC that V-59 activates, and the metabolic pathway it inhibits. The native function of Rv1625c signaling during infection is not established, and Rv1625c is the first AC that has been linked to a specific downstream metabolic pathway in Mtb (*23*). Our finding that the Rv1625c transmembrane domain is required for cholesterol ring catabolism expands on work by others showing that the catalytic domain is not the only functionally relevant component of this AC (*32, 33, 49*), but the mechanism that mediates its involvement in cholesterol catabolism remains to be determined. It is not known if the transmembrane domain of Rv1625c mediates protein-protein interactions that are required to complete cholesterol catabolism and whether this, or an alternative mechanism, is involved in maximizing regulation of cholesterol utilization by Rv1625c agonists. It is notable that the cholesterol utilization defect in the ΔRv1625 strain was likely limited to ring catabolism (Fig. 3a), and was not sufficient to impact bacterial growth in cholesterol media or macrophages (Fig. 1a, c), or to inhibit transcriptional activation of methylcitrate cycle genes by liberation of propionyl-CoA (Fig. 5 and Fig. 6a). Because degradation of the cholesterol side chain and rings can proceed in tandem, it is conceivable that a blockage in ring catabolism could be present in ΔRv1625 Mtb, while growth on cholesterol and liberation of propionyl-CoA and acetyl-CoA from the side chain are maintained. Two genes neighboring Rv1625c (*rv1626/pdtaR* and *rv1627c*) have been predicted to be required during cholesterol utilization (*43*). It will be helpful to determine whether Rv1625c interacts with these proteins, or others capable of regulating lipid metabolism in Mtb. Moreover, it would be interesting to examine whether the binding of V-59 to Rv1625c not only activates cAMP synthesis but also alters an interaction between Rv1625c and a relevant protein. This would help explain the distinct but overlapping cholesterol utilization defects we observed in WT Mtb treated with V-59, the ΔRv1625 strain, and TetOn-cAMP Mtb treated with different doses of Atc. The amount of cAMP produced by the TetOn-cAMP strain was most similar to V-59 treatment at the lower dose of Atc (50 ng/mL) tested (Fig. 4a). This dose was also correlated with less severe defects in growth in cholesterol media, ^14^CO_2_ release from [4-^14^C]-cholesterol, and propionyl-CoA liberation from cholesterol than the defects observed with V-59 treatment (Fig. 4b, c and Fig. 6b). Based on this and our observations in the ΔRv1625 strain, it is interesting to speculate that V-59 interacts with the Rv1625c protein, altering both a relevant protein interaction and increasing cAMP synthesis, both of which contribute to V-59’s total effect on cholesterol utilization. However, the mechanism and kinetics of inducing cAMP synthesis with the TetOn-cAMP system are distinct from V-59 treatment, limiting our ability to conclude that a particular dose of Atc mimics V-59 treatment. Future experiments to identify cholesterol degradation intermediates that accumulate in WT Mtb treated with V-59 could help clarify this by allowing comparison of step(s) of the cholesterol degradation pathway that are blocked by V-59 treatment versus Tet-On cAMP induction or loss of the Rv1625c protein alone.

These findings expand our limited understanding of how cAMP signaling can alter metabolism in Mtb, and it remains to be determined whether a downstream cAMP-binding protein is required for V-59 or TetOn-cAMP induction to inhibit cholesterol utilization in Mtb. We investigated Rv0998/Mt-Pat because it is a cAMP-binding lysine acetyltransferase that was previously shown to acetylate and inactive the acetyl-CoA/propionyl-CoA ligase (Rv3667/Acs) and various FadD enzymes in Mtb, which can regulate incorporation of 2- and 3-carbon precursors into central metabolism (*27, 28*). Mt-Pat was not required for V-59 to inhibit Mtb growth (Supplementary Fig. 5a) or induction of MCC genes (Supplementary Fig. 5b) in cholesterol media. This is consistent with data showing that acetate rescues growth during both V-59 treatment and TetOn-cAMP activation (Fig. 1b, Supplementary Fig. 5c), suggesting Acs has not been inactivated by Mt-Pat in either condition. The FadD enzyme Rv3515c/FadD19 initiates cholesterol side chain degradation (*6*) but FadD19 is not a confirmed target of inactivation by Mt-Pat, and other FadD enzymes that are known to be acetylated by Mt-Pat are not established steps in cholesterol breakdown (*28*).

Although Mt-Pat likely does not mediate inhibition of cholesterol utilization downstream of V-59 treatment or TetOn-cAMP induction, it is unclear whether any of the other eleven predicted cAMP-binding proteins in Mtb are involved in this mechanism (*23*). We examined the predicted operon of one other cAMP-binding protein, the transcription factor CRP_MT_, and found that a handful of these genes were differentially expressed in a cAMP-dependent and/or Rv1625c-dependent manner during growth in cholesterol media (Fig. 7). Notably, three of these genes are required for optimal growth of Mtb in cholesterol media and/or in mouse models of TB, but are not directly involved in cholesterol side chain or ring breakdown. Most of the remaining genes do not have established functions, making it difficult to predict how changes in their expression could impact specific aspects of Mtb physiology. Further experiments are needed to determine whether these transcriptional changes are indeed mediated through CRP_Mt_ activity, and whether this contributes significantly to the cholesterol utilization defects observed in Mtb during V-59 treatment or in the ΔRv1625c strain. Alternatively, because cholesterol uptake by the bacterium and initiation of cholesterol side chain breakdown by FadD19 are both ATP-dependent processes, it is possible that ATP depletion occurring during increased cAMP synthesis mediates inhibition of cholesterol utilization.

Compared to the previously-published Rv1625c agonist V-58 (*36*), V-59 represents a significant advance because it is the first Rv1625c agonist that is suitable for use in an *in vivo* model of TB. V-59 also provided a basis for understanding how to improve Rv1625c agonists through medicinal chemistry. We completed a medicinal chemistry effort focused on addressing the structural liabilities of V-59 and improving the potency of its activity against Mtb. This identified mCLB073, a stable molecule with improved potency and desirable chemical, pharmacological, pharmacokinetics and safety properties, which makes it a good drug candidate for clinical testing and a useful compound for further *in vivo* studies of this pathway in Mtb. While investigating the potency of compounds in this series with constrained piperidine rings, we also identified molecules whose potency differed widely based on the chirality of their azabicyclic rings. In the future, it would be interesting to identify the underlying explanation for the differential potency of these compounds on Rv1625c activity through structural biology.

Chemical tools like V-59/mCLB073 that can be used to modulate distinct pathways in Mtb in infection models are valuable because it is difficult to predict the *in vivo* metabolic state and vulnerabilities Mtb experiences during infection (*50–54*). We developed V-59/mCLB073 as the first compounds that can effectively modulate cAMP synthesis and block cholesterol utilization in Mtb, while being suitable for *in vivo* studies. Single-agent studies using V-59/mCLB073 established that cAMP induction modestly impairs bacterial growth and lung pathology in mouse models of TB (Fig. 8). A recent study also found that the “MKR superspreader” strain of Mtb, an emergent multidrug-resistant strain of the modern Beijing lineage, displayed enhanced upregulation of cholesterol utilization genes during macrophage infection relative to the H37Rv reference strain (*13*). V-59 was used to show that activating Rv1625c to inhibit cholesterol utilization in the MKR strain selectively reduced intracellular survival of the bacteria in infected macrophages (*13*). This suggests that efficacy of Rv1625c agonists may be potentiated by cholesterol-related metabolic adaptations that are especially crucial to the intracellular survival of at least one multi-drug resistant strain of Mtb. Identifying this link between cAMP signaling, cholesterol utilization, and Mtb fitness during infection is important because it is challenging to target metabolic pathways in Mtb that are not redundant and are sufficiently distinct from human pathways to limit side-effects (*27, 50, 55*). Numerous studies have suggested that cholesterol utilization is a key metabolic adaptation that supports Mtb survival during chronic infection (*7–11*), but the efficacy of single-step inhibitors of cholesterol degradation may be limited, unless they are able to cause accumulation of toxic metabolites in the bacterium (*6, 8-10, 56*). Inhibitors that block this pathway early and/or shut down multiple steps present one desirable alternative. This study revealed that the Rv1625c agonists V-59/mCLB073 are an improvement over the single-step cholesterol degradation inhibitors we reported previously; these compounds inhibit cholesterol catabolism early and/or at both side chain and ring degradation steps and display excellent pharmacokinetic properties. The optimized compound mCLB073 will facilitate future studies examining how Rv1625c agonists alter Mtb fitness in animal models that more faithfully recapitulate the complexities of the granuloma microenvironment present in human TB. Much remains to be discovered about the restrictions Mtb encounters *in vivo* when the bacterium co-catabolizes multiple complex substrates, and future infection studies that combine V-59/mCLB073 treatment with chemical modulators of other pathways in Mtb may provide interesting insights on this topic.

In summary, we have shown here that activating cAMP synthesis in Mtb, either by activating Rv1625c AC activity with a small molecule agonist or by inducing expression of the minimal catalytic subunit of Rv1264, blocks cholesterol degradation. Rv1625c is also the first AC in Mtb to be linked directly to a particular downstream pathway. Mtb has a cadre of structurally-diverse ACs and predicted cAMP-binding proteins with mostly uncharacterized functions, which may represent potential to alter pathways beyond cholesterol utilization in this bacterium. However, it is unknown whether agonists for ACs other than Rv1625c would have comparable downstream effects in Mtb. In other pathogenic bacteria, cAMP signaling is known not only for coordinating changes to carbon metabolism, but also for mediating diverse functions including biofilm formation, virulence gene expression, and secretion systems (*14*). This study identified AC agonists suitable for *in vivo* studies of cAMP signaling in Mtb, which revealed that chemically activating cAMP synthesis may be an untapped mechanism for manipulating bacterial fitness during infection, redefining the traditional mechanism for an anti-virulence compound. In Mtb, AC activation is able to stall at least one metabolic pathway that supports *in vivo* survival, with the potential to yield a new antibiotic. It is interesting to speculate whether additional AC agonists could be developed as tools to study cAMP signaling during infection, or as antibiotics, in other bacteria. The mechanism(s) by which inducing cAMP synthesis modulates cholesterol utilization in Mtb is not yet fully explained, and will be an important area for future studies as our knowledge of the role cAMP signaling plays in Mtb physiology continues to expand.

## Materials and Methods

### Bacterial culture

Unless noted, Mtb strains were grown in Middlebrook 7H9 medium supplemented with glycerol and OADC (oleic acid, albumin, dextrose, catalase) (Difco). 7H12 medium contained Middlebrook 7H9 powder (Becton Dickinson), 0.1% casitone, and 100 mmol MES free acid monohydrate, pH 6.6. Prior to culturing in media containing different carbon sources, bacteria were washed in 7H12 media without additional carbon sources. Cholesterol was added as tyloxapol:ethanol micelles to a final concentration of 100 µM (*29*). Where specified, 0.1% acetate was added. All liquid media contained 0.05% tyloxapol (Acros Organics). Mtb was cultured on Middlebrook 7H10 agar supplemented with glycerol and OADC (Difco). Strains were maintained with selective antibiotics as described in Table S2. Anhydrotetracyline was prepared in 100% EtOH and used at final concentrations of 500 ng/mL or 50 ng/mL.

### Construction of mutants and TetOn-cAMP strains

The *rv1625c* gene was disrupted in CDC1551 Mtb using allelic exchange (*38*). ΔRv1625c was complemented by overexpressing full-length Rv1625c or the transmembrane domain (amino acids V1-D204), from an integrating vector under the control of the *hsp60* promoter. The TetOn-cAMP strain expresses the His-tagged catalytic domain of Rv1264, under control of the *p606* Atc-inducible promoter. A single amino acid change (D265A) was introduced in the *rv1264* sequence using site-directed mutagenesis. Base pairs 794-795 were mutated (GAC to GCG) and confirmed by sequencing. Details of strains and constructs are listed in Table S2.

### Growth inhibition measurements

Growth assays were conducted by inoculating Mtb strains into liquid media at an OD_600_ of 0.05, and measuring the OD_600_ at three-day intervals. Compounds were added at the indicated concentrations initially and every three days throughout. For EC_50_ measurements, Mtb strains were pre-grown in 7H12+acetate media and assayed as described (*29*). For macrophage infections, bone marrow-derived macrophages (BMDMs) were isolated and differentiated from BALB/c mice (*38*). BMDMs were seeded in media without antibiotics in 24-well plates before infection. Cells were infected with Mtb. After 2 hours, extracellular bacteria were removed and replaced with fresh media containing indicated treatments. Media was replaced every 24 hours for the duration of the experiment. Fold changes in CFU’s were determined by lysing macrophages in SDS (0.01%) and plating on agar plates.

### ^14^CO_2_ release experiments

Catabolism of [4-^14^C]-cholesterol to ^14^CO_2_ was quantified as described previously, with minor modifications (*38*). Briefly, Mtb cultures were pre-grown in 7H12+cholesterol+acetate media for one week and adjusted to an OD_600_ of 0.5. DMSO or V-59 were added 45 minutes prior to adding [4-^14^C]-cholesterol. For TetOn-cAMP experiments, bacteria were inoculated at an OD_600_ of 0.1 into 7H12+cholesterol+acetate media and treated with EtOH, Atc, or V-59 at the indicated concentrations overnight, and again one hour prior to beginning the ^14^CO_2_ release assay. Cultures were adjusted to an OD_600_ of 0.5 in their respective media and [4-^14^C]-cholesterol was added. In both cases, ^14^CO_2_ released from the vented Mtb culture flasks was collected as described (*38*).

### Thin-layer chromatography

Mtb cultures were grown to an OD of 0.6, then inoculated at an OD_600_ of 0.4 into 7H12+cholesterol media for three days. Cultures were concentrated in their respective media, then [4-^14^C]-cholesterol was added, the culture supernatant was collected after 24 hours, extracted in ethyl acetate, quantified by scintillation counting, and equal counts (10,000 CPM per lane) were spotted for each sample on a silica gel TLC plate (EMD Chemicals). Plates were run in toluene:acetone (75:25, v/v) and imaged by phosphorimaging.

### Heterologous expression of Rv1625c in *cya*^-^ E. coli

The cya-deficient *E. coli* strain HS26, derived from the TP610 strain, was transformed with the pMBC530 plasmid expressing the full Rv1625c ORF or the empty vector control plasmid pMBC529 and the strains were grown and induced as previously described (*36*). After 18 hours, sample OD_600_ values were recorded and supernatants were collected and used to quantify cAMP by ELISA.

### Quantification of bacterial cAMP

Bacteria were pre-grown in 7H12+cholesterol+acetate before inoculation into fresh media at an OD_600_ of 0.1 containing either a cAMP-inducing compound (V-59 or Atc) or vehicle control. After 24-hour incubation, bacteria were collected by centrifugation, and the supernatant was reserved to measure secreted cAMP. The pellet was washed with fresh media, resuspended in lysis buffer (0.1M HCl, 1% Triton X-100 in H_2_O), and disrupted by bead beating (MP Biomedical). Cell-free lysates were reserved to measure internal cAMP by ELISA (Enzo Life Sciences). The sum of the internal and external values, or the external values alone, for each sample were used to estimate the total cAMP produced per 1×10^8^ bacteria.

### RNA-seq analysis

Bacteria were cultured in 7H9OADC before inoculation into fresh media containing a cAMP-inducing compound (10 µM V-59 or 500 ng/mL Atc) or vehicle control (DMSO or EtOH) at an OD_600_ of 0.1. The following day, bacteria were inoculated into 7H12+cholesterol media containing fresh compound or vehicle control. After four hours, cultures were pelleted by centrifugation, washed with guanidine thiocyanate-based buffer and stored at −80°C. Pellets were washed and suspended in Trizol LS (Ambion) and lysed by bead beating. Total RNA was isolated by chloroform extraction and precipitated in isopropanol with GlycoBlue reagent (Thermo Fisher).

RNA was resuspended in nuclease-free water, genomic DNA contamination was removed using the Turbo-DNA free kit (Invitrogen), and rRNA was depleted using the Ribo-Zero Gold rRNA removal kit (Illumina). Sample quality was determined via Fragment Analyzer (Advanced Analytical) and TruSeq-barcoded RNAseq libraries were generated with the NEBNext Ultra II Directional RNA Library Prep Kit (New England Biolabs). Sequencing was performed at the Cornell University Transcriptional Regulation and Expression Facility on a NextSeq500 instrument (Illumina) at a depth of 15 M single-end 75 bp reads. Reads were trimmed for low quality and adaptor sequences with TrimGalore, and aligned to the *Mycobacterium tuberculosis* CDC1551 reference genome (GCA_000008585.1) with STAR. DESeq2 was used with default parameters and alpha = 0.05 to generate the differential gene expression results. Multiple test correction was performed using the Benjamini Hochberg method.

### Propionyl-CoA reporter Assays

Fluorescent *prpD’*::GFP assays in liquid media were conducted in 7H12+cholesterol+acetate as described (*38*). BMDMs were infected at an MOI of 5. Extracellular Mtb was removed after 2 hours, and replaced with fresh media containing DMSO or V-59. For intracellular TetOn-cAMP experiments, the TetOn-cAMP *prpD’*::GFP strain was pre-induced in 7H9OADC media with EtOH, Atc (500 ng/mL), or V-59 (10 µM) for 24 hours prior to infection. After extracellular Mtb was removed, fresh media containing EtOH, Atc (3 µg/mL), or V-59 (10 µM) was added to the infected BMDMs. After 24 hours, infected BMDMs were scraped and fixed in paraformaldehyde. Fixed BMDMs were suspended in lysis buffer (0.1% SDS, 0.1 mg/mL Proteinase K in H_2_O) and lysed by passage through a 25-gauge needle. Pellets were retained for analysis. The GFP MFI was quantified from 10,000 bacteria by flow cytometry and analyzed using FlowJo (Becton Dickinson). Details of reporter plasmid constructs are listed in Table S2.

### Pharmacokinetic studies

Pharmacokinetic profiles were determined in male CD-1 mice after single dose oral administration (20 mg/kg). Stock solutions were prepared in 75% PEG 300, 25% D5W. An aliquot of the dose solutions was taken before and after dosing, and stored at –20°C for subsequent analysis. Blood samples were collected at standard time points through a retro-orbital bleed from 3 mice per time point. Heparinized blood was collected from each mouse and centrifuged to separate plasma for quantification of drug concentration by LC/MS analysis.

### Analysis of spontaneous resistance to mCLB073

The frequency of spontaneous resistance to mCLB073 *in vitro* was estimated by plating Mtb on cholesterol agar plates containing mCLB073 (25 µM to 100 µM). Cholesterol was dissolved in 500 mM methyl-β-cyclodextrin and added to 7H10 agar at 100 μM. 7×10^5^ CFU WT Erdman Mtb was spread per 150 mm plate and colonies were enumerated. Mutant clones were isolated, and subjected to EC_50_ assay. The *rv1625c* region was amplified by PCR and sequenced to determine the location of each mutation.

### Mouse infections

Animal work was approved by Cornell University IACUC (protocol number 2013-0030). All protocols conform to the USDA Animal Welfare Act, institutional policies on the care and humane treatment of animals, and other applicable laws and regulations. Isoflurane was delivered via nebulizer for anesthesia during oral delivery of compounds. Euthanasia was performed via delivery of carbon dioxide. Six to eight-week-old BALB/cJ mice (Jackson Laboratories) were infected with 1000 CFU of Erdman Mtb intranasally. C3HeB/FeJ (Kramnik) mice (Jackson Laboratories) were infected with 500 CFU of Erdman Mtb intranasally. At weeks 4 through 8 post-infection, compounds or vehicle controls were administered once-daily by oral gavage. For aerosol infection, BALB/cJ mice (Jackson Laboratories) were infected via an aerosol inhalation exposure system (Glass-Col) with a calibrated dose of 200 CFU of Erdman Mtb. Experimental compounds or vehicle control were administered once-daily at weeks 4 through 8 post-infection by oral gavage. After treatment, lung tissues were collected and processed for histology and CFU enumeration. For CFUs, lungs were homogenized in PBS, 0.05% Tween-80 and plated on 7H10 OADC. Inflammatory area was scored by measuring the percent of tissue inflamed (granulomatous tissue with peripheral and peribronchial lymphocytes and plasma cells) per low power microscopic field. In a blinded fashion, 4-15 fields were analyzed per lung sample.

## Statistical Analysis

Averages were chosen as a measure of central tendency throughout. Analyses were performed using GraphPad Prism. When data were expected to fit an approximately normal distribution Ordinary one-way ANOVA was applied. For multiple comparisons between pre-selected pairs of control versus treatment groups, Sidak’s multiple comparisons test was used. For comparisons between multiple treatment groups relative to a single control group, Dunnet’s multiple comparisons test was selected. Ordinary two-way ANOVA with Tukey’s multiple comparisons test was used when it was necessary to analyze the impact of both the strain background and compound versus control treatments. Data from mouse experiments were analyzed using non-parametric tests. For a single pre-planned comparison, a Mann-Whitney test was selected. For multiple comparisons, the Kruskal-Wallis test and Dunn’s multiple comparisons test were used. Differences with *P* values < 0.05 were considered significant. For RNA-seq results, significance of the log_2_ fold-change between groups was assigned based on the adjusted p-value for differential expression analysis, where significance was assigned when the p-adj value was < 0.05.

## Acknowledgements

We thank Kathleen McDonough for pMBC529, pMBC530, and the HS26 *E. coli*. We thank Dirk Schnappinger for the TetOn plasmids. We thank Véronique Dartois and Jansy Sarathy for completing the caseum binding assays. We thank Jen Grenier, Ann Tate, and Faraz Ahmed from Cornell TREx for assisting with RNA-seq sample analysis.

## Author contributions

K.W. constructed and characterized the Mtb strains, conducted the reporter assays, performed the RNA-seq experiments, did the radiolabeling experiments, and conducted the cAMP quantitation. C.M., isolated and characterized the spontaneous mutants. K.W., C.M., L.H., and B.V. conducted the in vivo modeling. A.W., B.Q., M.L., C.M., and H.M.P. conducted the chemical synthesis, activity characterization, and PK studies. T.S. conducted the pathology analysis. All authors contributed to the experimental design and data analysis. K.W., and B.V. wrote the manuscript, which was edited by all authors.

## Data and materials availability

All data supporting the findings of this study are presented in the article and its associated supplementary files. The complete RNA-seq dataset will be submitted to NCBI GenBank (accession number pending). Reagents can be requested and made available to users with verified approvals.

## List of Supplementary Materials

### Materials and Methods

#### Compound optimization

To identify mCLB073, the potency, safety, and pharmacokinetic properties of the oxadiazole series represented by V-59 was optimized. The cellular activity of V-59 analogs were assessed based on *in vitro* potency against Mtb in cholesterol-based media and intramacrophage assays, and cytotoxicity counterscreens against mammalian cells to ensure high selectivity.

#### Compound formulations for *in vivo* experiments

V-59 and isoniazid (Sigma Aldrich) were solubilized in 10% DMSO, 70% PEG 300, and 20% D5W (Dextrose 5% in ddH_2_O) with heating and sonication for intranasal BALB/c and C3HeB/FeJ experiments. For mCLB073 studies, compounds were solubilized at the indicated concentrations in 0.5% methyl cellulose, 0.5% Tween-80 (intranasal infection experiment) or 10% 2-hydroxypropyl-β-cyclodextrin + 10% lecithin (aerosol infection experiment) to obtain a fine suspension. Doses of 0.1mL were delivered by oral gavage.

#### In vitro cytotoxicity assay

In vitro cytotoxicity was evaluated in HepG2 cells (ATCC HB-8065, human liver) and in HEK293 cells (ATCC CRL-3216, human kidney). Cells were cultured in Dulbecco’s modified Eagle’s medium (DMEM) supplemented with 10% fetal bovine serum (FBS) and maintained in a humidified incubator (37°C in 5% CO_2_). Cells were harvested by trypsin treatment, collected by centrifugation, resuspended in assay medium with FBS (2 %) at the cell density of 7.5×10^4^ (HEK293T) or 5.0×10^4^ (HepG). Cell suspensions were dispensed (5 µL per well) into 1,536-well white microtiter plates pre-spotted with test compounds (Echo Labcyte), covered with metal lids (GNF), and incubated for 72 hours at 37°C. After 72 hours, CellTiterGlo (Promega) reagent was diluted 1:1 with ultra-pure water and dispensed into the plates (2 uL per well). Luminescence was measured with a microplate reader (EnVision), and the cytotoxic concentration, half maximal effect (CC_50_) was calculated for each compound with Genedata software. Test compounds were normalized to DMSO only wells, with puromycin (10 µM) as a positive control.

### Manual patch clamp assay for hERG (human ether-a-go-go-related gene) potassium channel inhibition

hERG inhibition was evaluated in CHO cells that stably express the hERG potassium channel (Sophion Biosciences). Cells were cultured in F12 Hams medium supplemented with 10% FBS, Hygromycin (100 µg/ml) (Thermo Fisher Scientific), and G418 (100 µg/ml) (Thermo Fisher Scientific) and maintained in a humidified incubator (37°C in 5% CO_2_). The test compounds were dissolved in DMSO (100 %) to obtain sub-stock solutions for different test concentrations and further diluted into media to achieve final concentrations for testing in cell culture. Final DMSO concentration did not exceed 0.30% in the compound treated, vehicle control, or positive (Amitriptyline) control.

The patch clamp assay was conducted with the extracellular solution (mM): NaCl 145, KCl 4, CaCl_2_ 2, MgCl_2_ 1, glucose 10, HEPES 10, pH = 7.4 and Osmolarity 290∼320 mOsm, and the intracellular solution (mM): KOH 31.25, KCl 120, EGTA 10, MgCl_2_ 1.75, CaCl_2_ 5.374, HEPES 10, Na-ATP 4, pH=7.2 and Osmolarity 280∼310 mOsm. Compounds were tested at room temperature using a USB amplifier (HEKA Elektronik). Output signals were low-pass filtered at 3 KHz and controlled with PatchMaster software (HEKA Elektronik). For quality control, the minimum seal resistance was set at 500 MOhms, and minimum specific hERG current (pre-compound) was 0.4 nA. Micropipettes (Harvard Apparatus) were pulled with a programmable micropipette puller (NARISHIGEPC-10Puller). The pipette tip resistance was between 2 ∼MOhms The recorded cells were continuously perfused with extracellular solution (Octaflow) at a flow rate of ∼1 ml/min mounted on the stage of an upright microscope (Nikon). The perfusion tip was manually positioned and five concentrations were tested in duplicate. Voltage command protocol: From the holding potential of −80 mV, the voltage was first stepped to 60 mV for 0.850 sec to open the hERG channels. Then the voltage was stepped down to −50 mV for 1.275 sec, generating a tail current, which was measured and collected for data analysis. Finally, the voltage was stepped back to the holding potential (−80 mV) and repeated every 15,000 msec. This command protocol was performed continuously during the test (vehicle control, test compounds). Compound application: During the initial recording period, the peak current amplitude was monitored until stable (< 5% change) for 5∼10 sweeps. Once stabilized, drug perfusion started with the lowest concentration and continued until the peak current stabilized across 5 sweeps, or 5 minutes if peak current did not change.

Data analysis was carried out using PatchMaster software (HEKA Elektronik) Excel 2013 (Microsoft) and Prism 5 (GraphPad). Normalized current values for each test compound concentration were calculated from recorded current responses: (peak current measured under compound perfusion /peak current measured with vehicle perfusion) ×100%. The percent inhibition values for each test article concentration were calculated from recorded current responses according to the formula: (1 - peak current measured under compound perfusion / peak current measured with vehicle perfusion)×100%. The final IC_50_ was determined with curve fitting. I/I control =Bottom + (Top-Bottom)/(1+10^((LogIC50-X)*Hill Slope)) where X is the logarithm of concentration, I/I control is the normalized peak current amplitude, Top is 1 and Bottom is equal to 0.

### Plasma protein binding assay

Plasma protein binding to compounds was quantified using human plasma (BioIVT) or plasma from CD-1 mice (Beijing Vital River Laboratory Animal Technology Co., Ltd) using a rapid equilibrium diffusion (RED) method. Equilibrium of free compound is achieved by the diffusion of the unbound compound across the (10-14 kDa MWCO) dialysis membrane in 96-well Micro-Equilibrium Dialysis Devices (HTDialysis). Briefly, sterile plasma containing test compounds was added to the matrix side of chamber in a commercial dialysis plate with reconstituted dialysis strips. Isotonic sodium phosphate buffer was added to the peripheral chamber of the dialysis plate and the unit was placed in a humidified incubator at 37 °C with 5% CO_2_ on a shaking platform at 100 rpm for 4 hours. Following dialysis, 50 μL aliquots of the samples were taken from the buffer side and the matrix side of the dialysis device and quenched with acetonitrile (100%) containing internal standards tolbutamide (200 ng/mL), labetalol (200 ng/mL) and metformin (50 ng/mL).

The concentration of free and bound test compound was determined by LC/MS analysis. The % Unbound, % Bound, and % Recovery values were calculated using the following equations: %Unbound=100 × F / T and %Bound = 100 - %Unbound where [F] is the analyte concentration or peak area ratio of analyte/internal standard in the buffer on the peripheral side of the membrane, [T] is the analyte concentration or peak area ratio of analyte/internal standard on the matrix side of the membrane, and [T0] is the analyte concentration or the peak area ratio of analyte/internal standard in the loading matrix sample at time zero.

### Caseum binding assay

Caseum binding was quantified using caseum harvested from rabbit Mtb granuloma lesions as described (*53*). Briefly, sterile caseum was spiked with test compounds and added to the matrix chamber of a commercial plate based micro-equilibrium dialysis device (Thermo Scientific Pierce RED Device). Isotonic sodium phosphate buffer was added to the peripheral chamber of the RED device and the plate was incubated at 37°C for 4 hours. Equilibrium of free compound is achieved by the diffusion of the unbound compound across the (8 kDa MWCO) dialysis membrane. Following incubation, aliquots of the isotonic buffer and the plasma were taken at pre-determined time points and the concentration of free and bound test compound was determined by LC/MS analysis.

### CYP Inhibition evaluation in human liver microsomes

Inhibition of five separate P450 monooxygenases was quantified using human liver microsomes (Corning). Briefly, microsomes were prepared in potassium phosphate buffer (100 mM) pH 7.2 containing phenacetin (10 µM), diclofenac (5 µM), S-mephenytoin (30 µM), dextromethorphan (5 µM), and midazolam (2 µM) which are substrates for the 1A2, 2C9, 2C19, 2D6, and 3A4 monooxygenase enzymes, respectively. Human liver microsomes were loaded into wells of an incubation plate and prewarmed to 37°C. NADPH was added to the final concentration of 10 mmol to all wells and the plate was incubated for 10 minutes at 37°C. The reactions were terminated by adding acetonitrile (100%), containing the internal standards tolbutamide (200 ng/mL) and labetalol (200 ng/mL). Protein was removed from the samples by centrifugation and the P450 monooxygenase enzymatic products were quantified by LC/MS/MS analysis. The concentration of the enzymatic products in the reaction mixture was used to calculate the percent of inhibition vehicle control versus the test compound concentrations (0, 0.05, 0.15, 0.5, 1.5, 5.0, 15.0 and 50 µM). Data was fit using a non-linear regression analysis and the IC_50_ values were determined using 3- or 4-parameter logistic equation. IC_50_ values were reported as “>50 µM” when % inhibition at the highest concentration (50 µM) is less than 50%.

Equation for three parameters logistic sigmoidal curve:

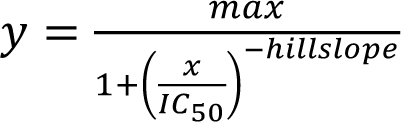

Equation for four parameters logistic sigmoidal curve:

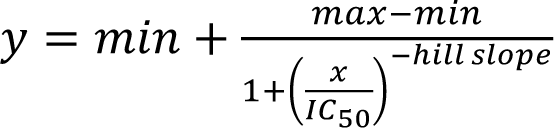

### Liver microsomal stability

Liver microsomes from mouse (Xenotech), human (Corning) rat (Xenotech) and dog (Xenotech) were prepared in potassium phosphate buffer (100 mM) pH 7.2. Experimental compounds were dissolved in DMSO (100%) to a stock concentration of 10 mmol. A working compound solution was made by diluting 5 μL of compound stock into 495 μL of acetonitrile (100%) to the final concentration of compound (100 μM) and acetonitrile (99%). Similarly, positive control compounds (Testosterone, Dichlofenac, and Propafenone) were dissolved in DMSO (100% to a stock concentration of 10 mM in DMSO, and were diluted with 495 μL of acetonitrile (100%) to the final concentration of compound (100 μM) and acetonitrile (99%). NADPH•4Na was dissolved into a 10 mM MgCl_2_ solution (10 unit/mL) generating as a 10x cofactor stock. Pre-warmed incubation plates were filled with 445 µL potassium phosphate buffer and incubated for 10 min at 37°C with constant shaking. Liver microsomes (54 µL) liver and NADPH stock (6 µL) were added to the plates. Working solution of compounds (5 µL) was added into the incubation plates containing microsomes and mixed thoroughly before incubating at 37°C for 60 min under constant shaking at 100 rpm. Reactions were stopped by the addition of 100 µl cold acetonitrile (100%) containing the internal standards tolbutamide (200 ng/mL) and labetalol (200 ng/mL). All plates were shaken for 10 min, then centrifuged at 4000 rpm for 20 minutes at 4°C. The cell free supernatant was transferred into HPLC water and mixed by shaking the plate for 10 min minutes prior to metabolite quantification by LC-MS/MS analysis.

The equation of first order kinetics was used to calculate T_1/2_ and CL_int(mic)_ (μL/min/mg):

Equation of first order kinetics:

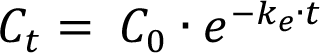

when 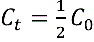

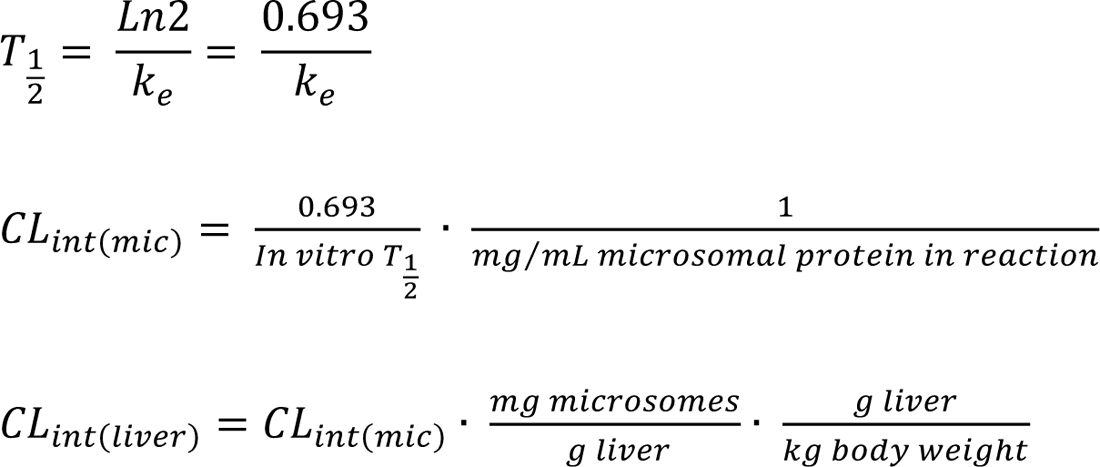

T_1/2_ is half life and CL_int(mic)_ is the intrinsic clearance

CL_int(mic)_ = 0.693/half life/mg microsome protein per mL

CL_int(liver)_ = CL_int(mic)_ * mg microsomal protein/g liver weight * g liver weight/kg body weight

Extraction ratio calculation

Extraction ratio (ER) = CLH/QH= CLint(liver)/(CLint(liver) + Qh)

Qh is Hepatic Blood Flow (mL/min/kg)

### Solubility

Stock solutions containing compound of interest (10 mM) were prepared in DMSO (100%) and diluted to a final concentration (0.2 mM) in phosphate buffer solution (50 mM) pH 6.8. The sample was loaded in Mini-UniPrep vials containing 0.2 µm filters (GE Halthcare), vortexed for 2 mins and shaken for 24 hours at room temperature. The Mini-UniPrep was compressed and the resulting filtrates were analyzed by HPLC/UPLC system to calculate the concentration with standard curve (1, 20, and 200 µM).

### Time over EC_50_ determination

The pharmacokinetic profile of mCLB073 was determined in male CD-1 mice after single dose oral administration (20 mg/kg). A 4 mg/ml stock solution of mCLB073 was prepared in 75% PEG 300, 25% D5W with vortexing to dissolve into solution. An aliquot of the dose solutions was taken before the dosing and after dosing, and stored at approximately –20°C for subsequent analysis. Blood samples were collected at standard time points (0.25h, 1h, 3h, 5h, 8h, 24h) through a capillary via a retro orbital bleed. At each time point, approximately 100μL of blood samples were collected from 3 mice per group. The blood was heparin treated and within 1 hour of collection the blood samples were centrifuged at 2500x g for 15 minutes at 4°C to collect the plasma for quantification of drug concentration via LC/MS. Time over EC_50_ was calculated by multiple EC_50_ with molecular weight. The equation is: EC_50_ line (ng/ml) = molecular weight (g/mol) × EC_50_ (µM, µmol/L).

### Western blot analysis

To confirm expression of His-tagged Rv1264 and Rv1264_D265A_ in the Tet-On Mtb strains, bacteria were grown to an OD of 0.6 in 7H9OADC and then inoculated into 7H12+acetate containing EtOH or Atc (500 ng/mL) at an OD of 0.15. The following day, an equivalent number of bacteria were pelleted for each strain and treatment, and the pellets were fixed in paraformaldehyde (4%) for one hour and stored at −80°C. Fixed Mtb pellets were suspended in SDS (1%) and probe sonicated (3 x 1 minute cycles). Bacterial debris was removed by centrifugation, and the supernatants were suspended in SDS PAGE loading buffer and boiled for 30 minutes with periodic vortexing.

Equivalent volumes of each sample were resolved by SDS-PAGE gel and transferred to nitrocellulose membranes. Western blotting was performed using either mouse anti-5His (Qiagen), or mouse anti-Mycobacterium tuberculosis GroEL2 (BEI Resources, Clone IT-70) primary antibody, and HRP-conjugated goat anti-mouse IgG (Jackson ImmunoResearch) secondary antibody. Chemiluminescent substrate (Thermo Fisher Scientific) was added and Westerns were imaged by film for anti-His or by ChemiDoc (Bio-Rad) for anti-GroEL2.

### cAMP production by cultured human cells

cAMP production was evaluated in HepG2 cells (ATCC HB-8065, human liver), HEK293 cells (ATCC CRL-3216, human kidney) and human monocyte derived macrophages (HMDM). HepG2 and HEK293 cells were cultured in DMEM supplemented with 10% FBS and maintained in a humidified incubator (37°C in 5% CO2). HMDM cells derived from human peripheral blood mononuclear cells (PBMCs) that were obtained from Elutriation Core Facility, University of Nebraska Medical Center. HMDM cells were cultured in DMEM supplemented with 10% human serum, L-glutamine (2 mmol), sodium pyruvate (1 mmol), penicillin (100 U/mL), and streptomycin (100 μg/mL) (Corning) and maintained in a humidified incubator (37°C in 5% CO_2_). Cells were cultured at a density of 1e6 per T-25 flask and were treated with V-59 (10 µM) for 24 hours before harvesting the cells with trypsin treatment (HepG2 and HEK293) or scraping into cold PBS (HMDM). Cells were harvested by centrifugation and cAMP levels were quantified from the cell pellet following, resuspension in lysis buffer (0.1M HCl, 1% Triton X-100 in ddH_2_O). The cell-free lysates were used to measure internal cAMP by ELISA (Enzo Life Sciences).

## Supplementary Figures

**Fig. S1.**
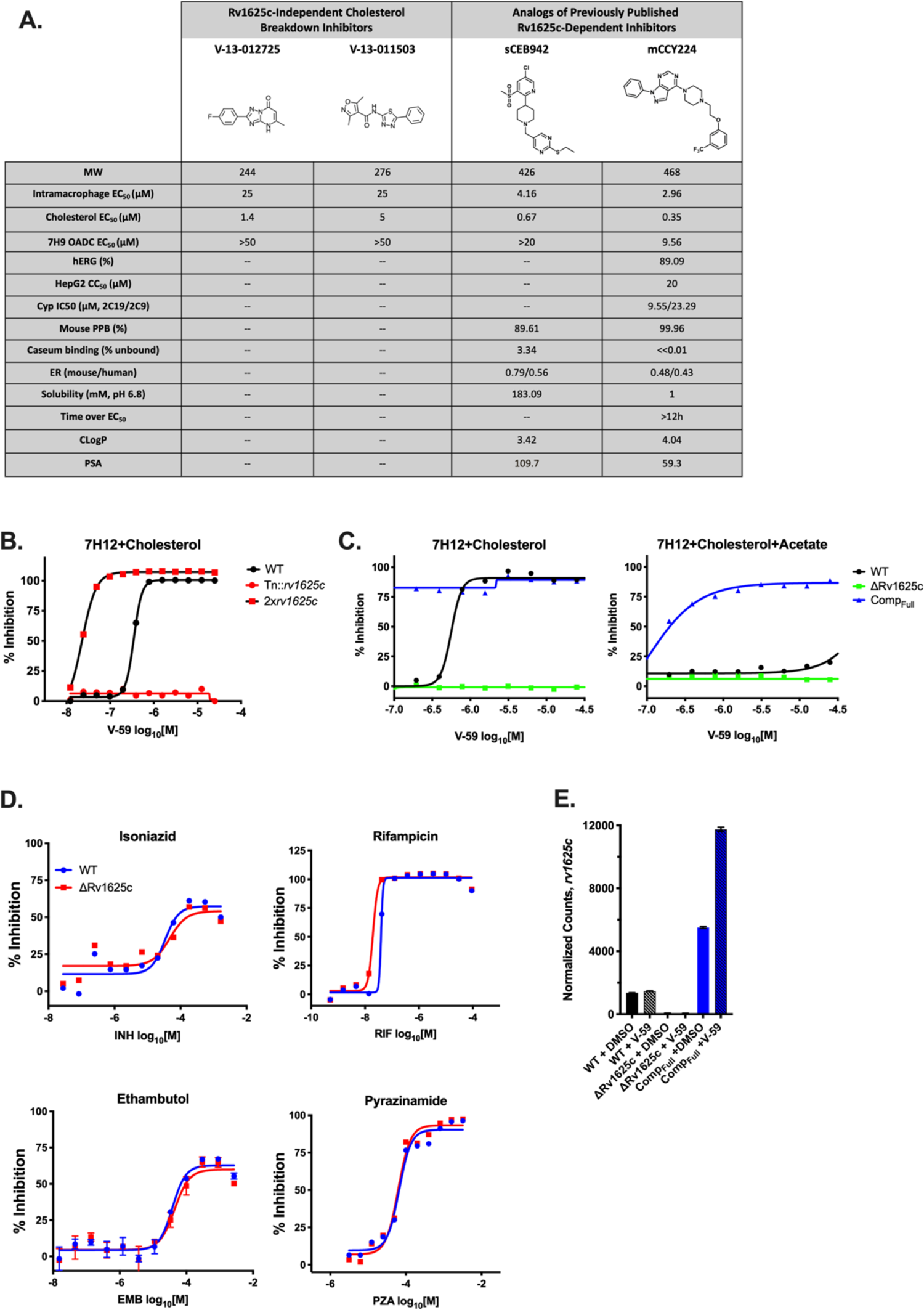
V-59 is structurally distinct from other cholesterol utilization inhibitors and the inhibitory activity of V-59 in cholesterol media is dependent on Rv1625c. (**A**) Chemical structure of previously published single-step cholesterol breakdown inhibitors, and resynthesized analogs of previously described Rv1625c-depedent inhibitors. (**B**) Inhibitory activity of V-59 against WT, the Rv1625c transposon mutant (Tn::*rv1625c*), and WT transformed with an overexpression plasmid expressing the *rv1625c* gene (2x*rv1625c*) in 7H12+cholesterol media. (**C**) Inhibitory activity of V-59 against WT, ΔRv1625c, and Comp_full_ in 7H12+cholesterol media (left) or 7H12+cholesterol+acetate media (right). Data shown are representative, from one experiment with two technical replicates. Symbols are mean data points, and curves display nonlinear fit of dose-response. (**D**) Effect of frontline antibiotics on WT and ΔRv1625 in 7H12+cholesterol (INH, RIF, EMB) or in MES-buffered 7H9OADC+glycerol, pH 5.9 (PZA). Data shown are representative, from one experiment with two technical replicates. Symbols are mean data points, and curves display nonlinear fit of dose-response. (**E**) RNA-seq derived normalized counts of *rv1625c* reads in WT, ΔRv1625c, and Comp_Full_ strains in 7H12+cholesterol media.

**Fig. S2.**
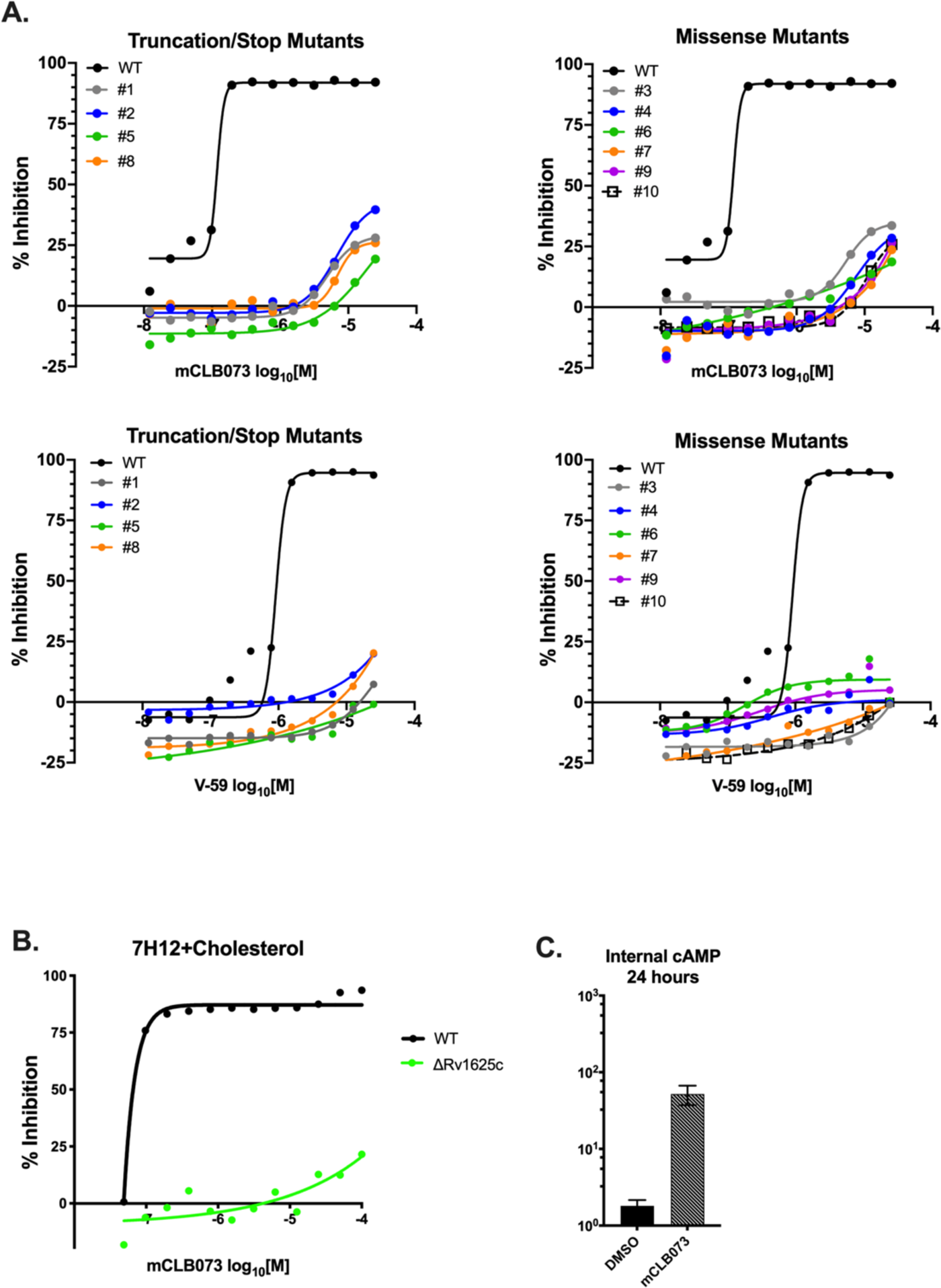
The activity of mCLB073 is dependent on Rv1625c. (**A**) Inhibitory activity of mCLB073 or V-59 against spontaneous resistant mutants that were generated during culture with mCLB073. (**B**) Inhibitory activity of mCLB073 against WT or ΔRv1625c in 7H12+cholesterol media. Data shown are representative, from one experiment with two technical replicates. Symbols are mean data points, and curves display nonlinear fit of dose-response. (**C**) Impact of mCLB073 on cAMP production in WT Mtb. ELISA was used to quantify cAMP from lysed cells 24 hours after addition of mCLB073, or vehicle control (DMSO). Data is from one experiment, with two technical replicates. Data are shown as means ± SEM.

**Fig. S3.**
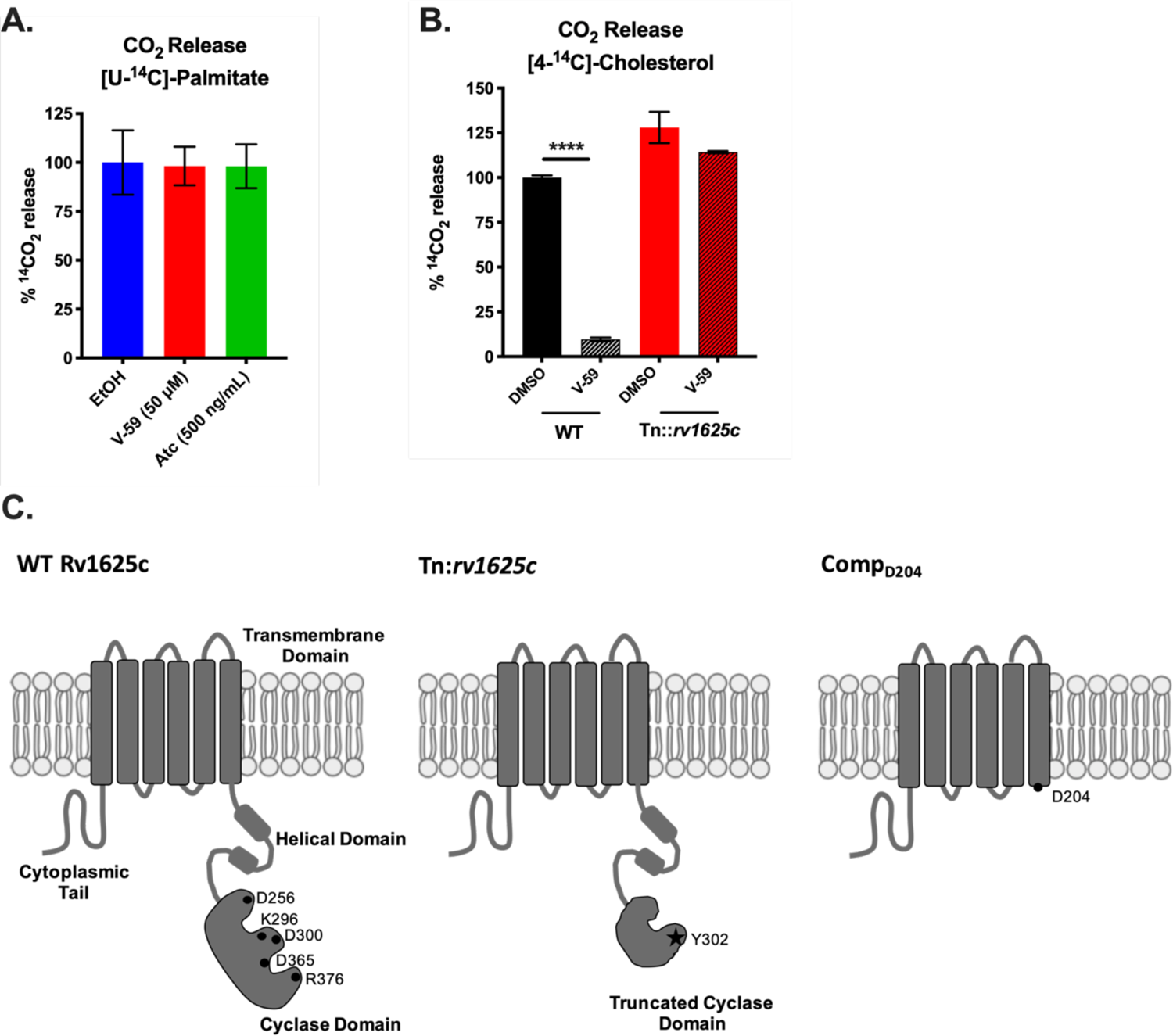
An intact cyclase domain of Rv1625c is required for V-59 to inhibit cholesterol degradation. (**A**) Catabolic release of ^14^CO_2_ from [U-^14^C]-palmitate in media containing fatty acid. EtOH (control), V-59, or Atc were added to the TetOn-cAMP Mtb cultures 24 hours prior to the start of the experiment. Data are from one experiment with three technical replicates, normalized to OD and quantified relative to EtOH control. Data are shown as means ± SEM (**B**) Catabolic release of ^14^CO_2_ from [4-^14^C]-cholesterol in media containing cholesterol and acetate. V-59 (10 μM) was added to the cultures once at the beginning of the experiments and DMSO was used as a vehicle control. Data are from one experiment with three technical replicates, normalized to OD and quantified relative to WT treated with DMSO. Data are shown as means ± SEM (\*\*\*\**P* < 0.001, One-way ANOVA with Sidak’s multiple comparisons test). (**C**) Schematic illustrating the topology of the N-terminal transmembrane domain and essential residues of the C-terminal cyclase domain of Rv1625c (left). Schematics illustrating modified Rv1625c constructs used in these studies (center and right).

**Fig. S4.**
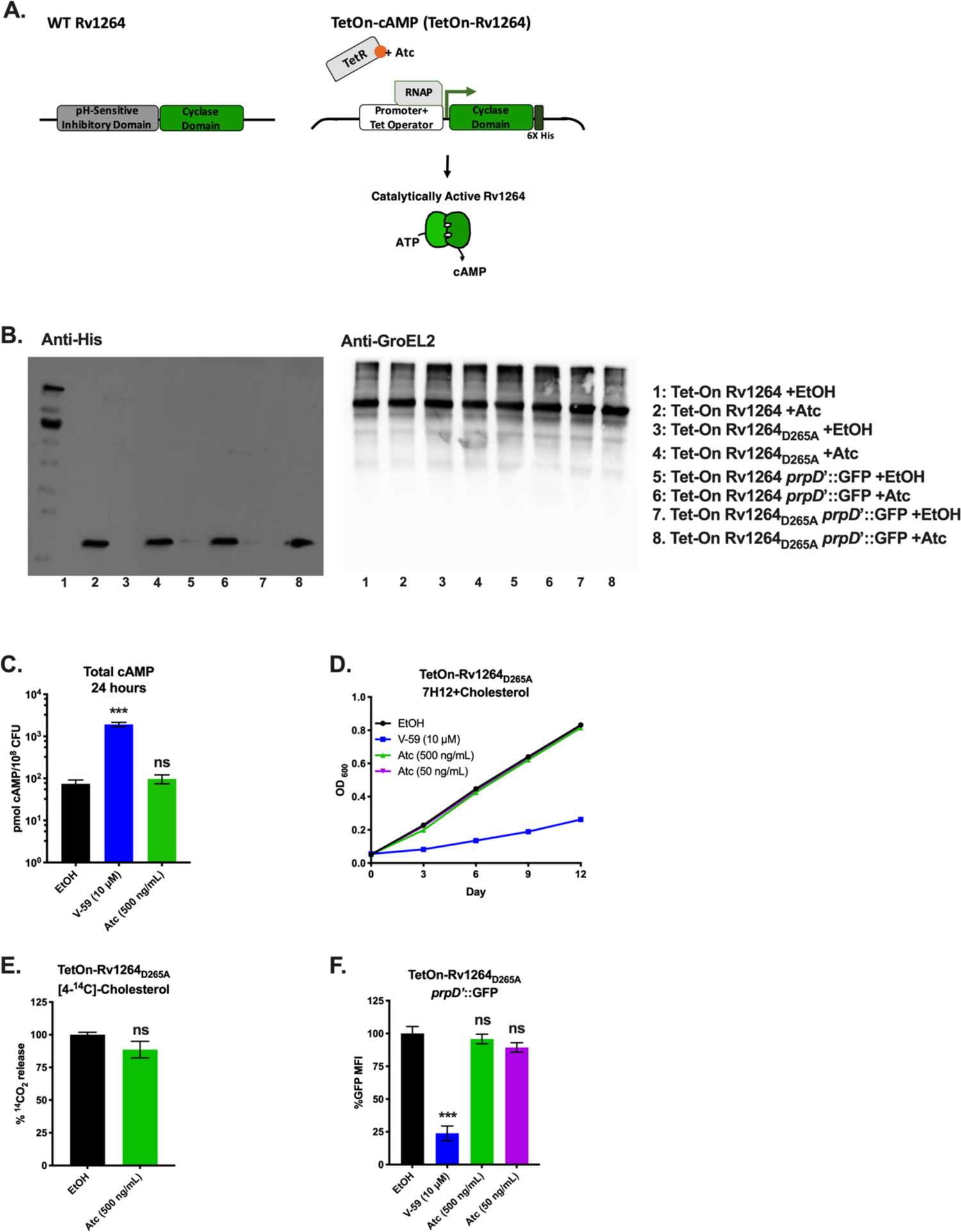
Construction and validation of TetOn-cAMP constructs. (**A**) Schematic illustrating the domains of the native Rv1264 adenylyl cyclase (left) and the design of the TetOn-cAMP construct (right). The TetOn-cAMP construct contains the minimum-necessary cyclase domain of Rv1264, and lacks the pH-sensitive inhibitory domain of the native Rv1264 protein. Expression of the cyclase domain is under control of a TetOn promoter. Upon treatment with Atc, release of the tetracycline repressor (TetR) causes initiation of transcription of the Rv1264 catalytic domain. The C-terminal end of the cyclase domain is His-tagged to allow immunoblotting. (**B**) Immunoblots of bacterial lysates confirm that the TetOn constructs are expressed in the presence of Atc and not the vehicle control EtOH. The anti-His blot detects the Rv1264 cyclase domain and the anti-GroEL2 blot is the loading control. (**C**) Impact of inducing the TetOn-Rv1264_D265A_ construct in media containing cholesterol and acetate. Cultures were treated with one dose of EtOH, V-59 (10 μΜ), or Atc at the indicated concentrations and samples were collected 24 hours later for ELISA. Data are displayed as total cAMP per 10^8^ Mtb. Data are from two independent experiments with two technical replicates each, shown as means ± SEM (\*\*\**P* < 0.001, One-way ANOVA with Dunnett’s multiple comparisons test). (**D**) Effect of inducing TetOn-Rv1264_D265A_ on the growth of Mtb in 7H12+cholesterol media, monitored by serial OD measurements. EtOH, V-59 (10 μM), or Atc at the indicated concentrations were added initially and every three days for the duration of the experiment. Data are from one experiment with three technical replicates, shown as means ± SEM. (**E**) Catabolic release of ^14^CO_2_ from [4-^14^C]-cholesterol in media containing cholesterol and acetate. The TetOn-Rv1264_D265A_ strain was treated with EtOH, V-59 (10 μM), or Atc at the indicated concentrations overnight and again one hour prior to the beginning of the experiments. Data are from two independent experiments with three technical replicates each, normalized to OD and quantified relative to EtOH vehicle control. Shown as means ± SEM (not significant, Student’s *t* test). (**F**) Relative GFP signal from the *prpD’*::GFP reporter in response to inducing TetOn-Rv1264_D265A_ in media containing cholesterol and acetate. Cultures were treated with EtOH, V-59 (10 μΜ), or Atc at the indicated concentrations. Data are normalized to EtOH vehicle control (\*\*\**P* < 0.001, One-way ANOVA with Dunnett’s multiple comparisons test). GFP MFI was quantified from 10,000 mCherry^+^ Mtb. Data are from two independent experiments with two technical replicates each, shown as means ± SEM.

**Fig. S5.**
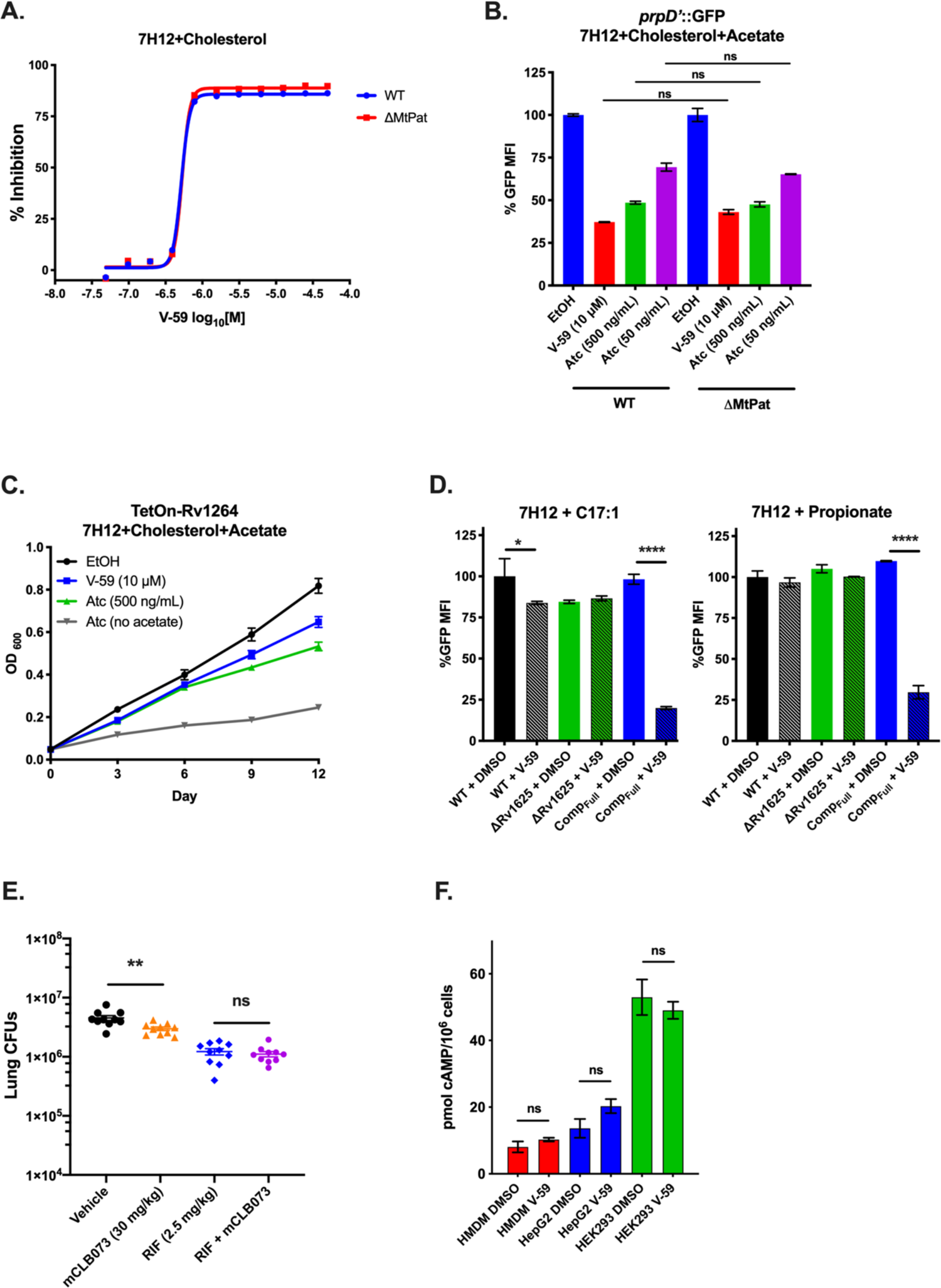
Activating cAMP synthesis inhibits lipid metabolism in an Mt-Pat independent mechanism, and can inhibit fatty acid utilization without increasing antibiotic tolerance or increasing mammalian cAMP synthesis. (**A**) Inhibitory activity of V-59 against WT or ΔMt-Pat in 7H12+cholesterol media. Data shown are representative, from one experiment with two technical replicates. Symbols are mean data points, and curves display nonlinear fit of dose-response. (**B**) Relative GFP signal from the *prpD’*::GFP reporter in WT versus ΔMt-Pat strains carrying the TetOn-cAMP construct in media containing cholesterol and acetate. Cultures were treated in parallel with EtOH, V-59 (10 μΜ), or Atc at the indicated concentrations. Data are normalized to EtOH vehicle control in each strain. Responses to cAMP inducing compounds were comparable between strains (not significant, Two-way ANOVA with Tukey’s multiple comparisons test). GFP MFI was quantified from 10,000 mCherry^+^ Mtb. Data are from one experiment with two technical replicates, shown as means ± SEM. (**C**) Effect of inducing TetOn-cAMP on the growth of Mtb in 7H12+cholesterol+acetate media. EtOH, V-59 (10 μM), or Atc at the indicated concentrations were added initially and every three days for the duration of the experiment. Data are from one experiment with three technical replicates, shown as means ± SEM. (**D**) Relative GFP signal from the *prpD’*::GFP reporter in Mtb treated with V-59 or DMSO, in 7H12 media supplemented with C17:1 or propionate. Data shown normalized to WT+DMSO. GFP MFI was quantified from 10,000 mCherry positive Mtb. Data are from one experiment with two technical replicates, shown as means ± SEM (\**P* < 0.05, \*\*\*\**P* < 0.0001, One-way ANOVA with Sidak’s multiple comparisons test). (**E**) Effect of mCLB073 treatment combined with a sub-optimal dose of rifampicin (RIF) in infected BALB/c mice. Data are from one experiment with 10 mice per group (\*\**P* < 0.01, Mann-Whitney test). Infections with Mtb in (A and B) were by the intranasal route. Infections with Mtb in (C and D) were by aerosol. All data are shown as means ± SEM. **(F)** Quantification of cAMP in human cell lines treated with V-59 (10 µM) or DMSO control. Data are from one experiment with two technical replicates, and are shown as means ± SEM. V-59 did not significantly increase cAMP (not significant, One-way ANOVA with Sidak’s multiple comparisons test).

**Table S1.**
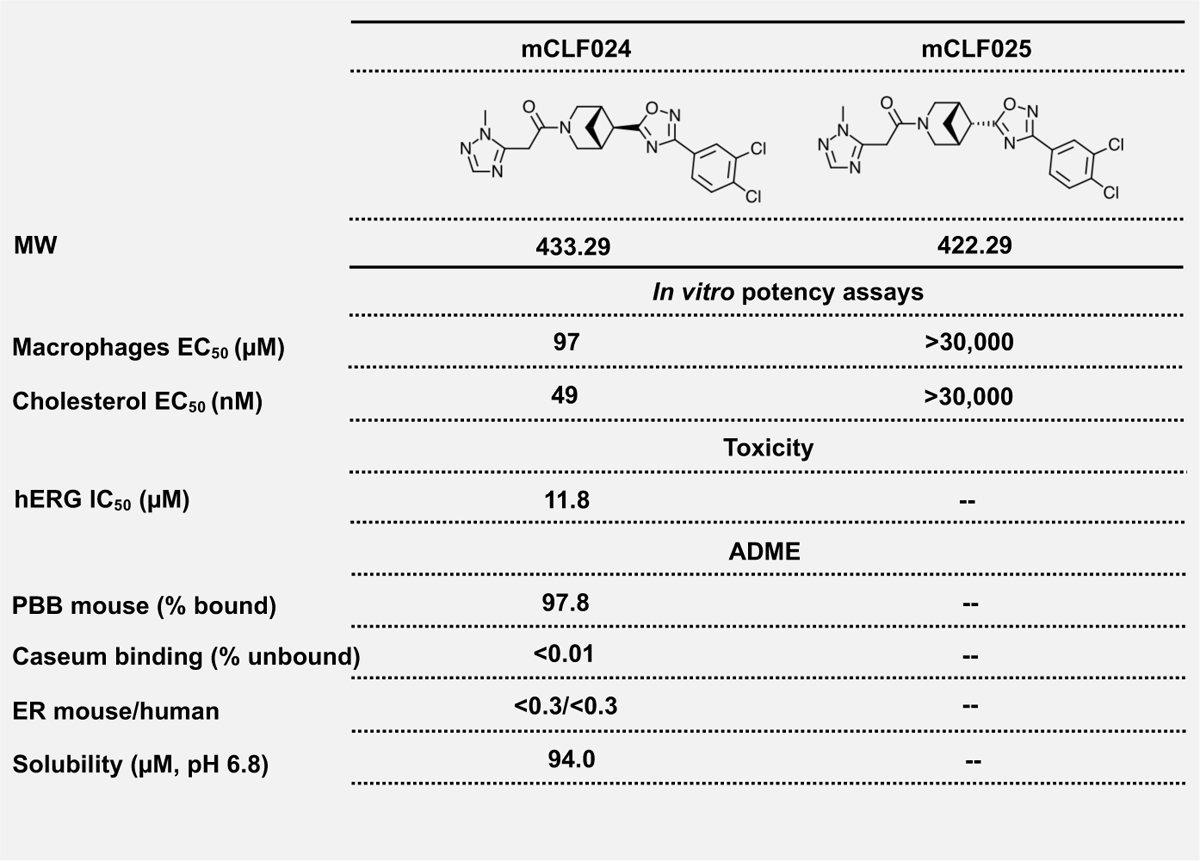
Structures and activities of isomer compounds. MW, molecular weight; --, not determined; EC_50_, half-maximal effective concentration; IC_50_, half-maximal inhibitory concentration; ADME, absorption, distribution, metabolism, excretion; PPB, plasma protein binding; ER, extraction ratio; PO, per os.

|  | mCLF024 | mCLF025 |
| --- | --- | --- |
| MW | 433.29 | 422.29 |
| *In vitro* potency assays |  |  |
| Macrophages EC<sub>50</sub> (μM) | 97 | >30,000 |
| Cholesterol EC<sub>50</sub> (nM) | 49 | >30,000 |
| Toxicity |  |  |
| hERG IC<sub>50</sub> (μM) | 11.8 | -- |
| ADME |  |  |
| PBB mouse (% bound) | 97.8 | -- |
| Caseum binding (% unbound) | <0.01 | -- |
| ER mouse/human | <0.3/<0.3 | -- |
| Solubility (μM, pH 6.8) | 94.0 | -- |

**Table S2.** Mtb strains used in these experiments

| Strain | Location of antibiotic resistance cassette insertion | Background | Antibiotic Resistance |
| --- | --- | --- | --- |
| Tn::Rv1625c | Mycomar transposon insertion at 904bp of *Rv1625c* | M. tuberculosis CDC1551 | Kanamycin |
| ΔRv1625c | (92-1256bp) of *Rv1625c* | M. tuberculosis CDC1551 | Hygromycin |
| ΔMt-Pat | (21-981bp) of *Rv0998* | M. tuberculosis CDC1551 | Hygromycin |

**Table S2. Mtb strains used in these experiments.**
| Strain | Plasmid Constructs | Background | Antibiotic Resistance |
| --- | --- | --- | --- |
| 2xRv1625* | pMV306+aacC4 (apramycin resistance)+<br>hsp60':Rv1625c (1-1329bp) | WT M. tuberculosis CDC1551 | Apramycin |
| Comp<sub>Full</sub>* | pMV306 + aacC4 (apramycin resistance) +<br>hsp60':Rv1625c (1-1329bp) | ΔRv1625c | Hygromycin, Apramycin |
| Comp<sub>D204</sub>* | pMV306+aacC4 (apramycin resistance) +<br>hsp60':Rv1625c (1-612bp) | ΔRv1625c | Hygromycin, Apramycin |
| TetOn-cAMP** | pDE1264 (pDO23A+Rv1264 (627-1191bp) ) +<br>pEN41A-T10M + pEN12A-p606 + pDE43-MEH | WT M. tuberculosis CDC1551 | Hygromycin |
| TetOn-Rv1264<sub>D265A</sub>** | pDE1264<sub>D265A</sub> (pDO23A+Rv1264<sub>D265A</sub>(624-1191bp) )+<br>pEN41A-T10M + pEN12A-p606 + pDE43-MEH | WT M. tuberculosis CDC1551 | Hygromycin |
| prpD':GFP | pCherry3 + kanamycin resistance +<br>prpD':GFP + smyc':mCherry | WT M. tuberculosis CDC1551 | Kanamycin |
| ΔRv1625c prpD':GFP | pCherry3 + kanamycin resistance +<br>prpD':GFP + smyc':mCherry | ΔRv1625c | Kanamycin |
| Comp<sub>Full</sub> prpD':GFP | pCherry3 + zeocin resistance +<br>prpD':GFP + smyc':mCherry | Comp<sub>Full</sub> | Hygromycin, Apramycin, Zeocin |
| Comp<sub>D204</sub> prpD':GFP | pCherry3 + zeocin resistance +<br>prpD':GFP + smyc':mCherry | Comp<sub>D265A</sub> | Hygromycin, Apramycin, Zeocin |
| TetOn-cAMP prpD':GFP | TetOn-cAMP + prpD':GFP + smyc':mCherry | WT M. tuberculosis CDC1551 | Hygromycin |
| ΔMt-Pat TetOn-cAMP prpD':GFP | TetOn-cAMP + kanamycin resistance +<br>prpD':GFP + smyc':mCherry | ΔMt-Pat | Hygromycin, Kanamycin |
| TetOn-1264<sub>D265A</sub> prpD':GFP | TetOn-Rv1264<sub>D265A</sub> + prpD':GFP + smyc':mCherry | WT M. tuberculosis CDC1551 | Hygromycin |

\*Apramycin gene sourced from Consaul SA, Pavelka MS Jr. Use of a novel allele of the Escherichia coli aacC4 aminoglycoside resistance gene as a genetic marker in mycobacteria. FEMS Microbiol Lett. 2004 May 15;234(2):297-301. doi: 10.1016/j.femsle.2004.03.041. PMID: 15135536.
\*\*Plasmids for Gateway cloning to construct TetOn-cAMP and TetOn-Rv1264<sub>D265A</sub> developed from Klotzsche M, Ehrt S, Schnappinger D. Improved tetracycline repressors for gene silencing in mycobacteria. Nucleic Acids Res. 2009 Apr;37(6):1778-88. doi: 10.1093/nar/gkp015. PMID: 19174563.

